# Structural enrichment for captive macaques - a systematic review and meta-analysis of behavioral outcomes

**DOI:** 10.1101/2025.09.09.675079

**Authors:** Jenny P. Berrio Sanchez, Christopher Musembi Makau, Dana Pfefferle, Sarah Neal, Otto Kalliokoski

**Affiliations:** Department of Veterinary and Animal Sciences. Section for Biomedicine. Faculty of Health and Medical sciences. University of Copenhagen. Ridebanevej 9, Frederiksberg. DK-1870. Denmark; Department of Veterinary Anatomy and Physiology, Faculty of Veterinary Medicine, University of Nairobi. P.O Box 30197-00100, Nairobi, Kenya; Welfare and Cognition Group, Cognitive Neuroscience Laboratory, German Primate Center, Kellnerweg 7, 37077 Goettingen, Germany; Department of Comparative Medicine, Michale E. Keeling Center for Comparative Medicine and Research. University of Texas MD Anderson Cancer Center.

**Keywords:** structural enrichment, macaque, welfare, behavior, housing

## Abstract

Macaques play an important role in biomedical research due to their genetic and physiological similarities to humans. However, their use raises significant ethical concerns regarding welfare, particularly in laboratory environments often lacking natural complexity. Environmental enrichment aims to promote welfare by encouraging natural behaviors and providing choice. This systematic review and meta-analysis synthesizes 61 years of research assessing the impact of structural modifications to the home cage (structural enrichment) on the behavior of captive macaques. We employed multilevel meta-analytical models to pool our data. Our findings demonstrate a net beneficial effect of structural enrichment on macaque behavior, notably in significantly reducing negative behaviors such as stereotypies, self-directed harm, and aggression. This positive effect was evident even when enrichment was added to already enriched conditions. We found that enrichment that incorporates varied physical and feeding strategies were most effective, underscoring that complex, varied environmental elements are crucial for welfare improvement. Simple increases in unfurnished space, by contrast, showed minimal benefit. The promotion of positive, species-typical behaviors was less consistent across studies, but nonetheless highlights structural enrichment’s potential to foster a broader natural behavioral repertoire. We also observed considerable variability in outcomes, suggesting that enrichment efficacy is influenced by multiple factors, and requires careful contextual assessment, as a ‘one-size-fits-all’ approach is rarely optimal.

## Introduction

Macaques play an important role in biomedical research given their genetic proximity to humans and similar physiology ^1^. These resemblances make them ideal for studying a wide array of human conditions and for assessing the safety and efficiency of new interventions. Compared to other non-human primates (NHPs), macaques are also more practical for research environments, largely because their small size makes housing and breeding easier^2^. However, their use in research is a subject of intense ethical scrutiny given the high capacity for sentience and suffering^3^. In terms of welfare, a major factor influencing the wellbeing of captive macaques is the housing environment where they spend most, if not all, of their lives. Lab primates often live in environments much simpler than their natural habitats. Such lack of stimulation can lead to distress^4^. Consequently, improving the macaques’ housing conditions has become an important goal among the research community. This is not just because of moral obligation^3^; poor welfare can also negatively affect the validity and reproducibility of research findings^5,6^.

The housing environment for captive macaques should ideally provide enough space and stimulation to allow for the expression of their full range of species-typical behaviors^7^. Macaques are highly social animals with complex filial and hierarchical structures^8^. When they are not interacting socially, they primarily spend their time foraging and interacting with their surroundings^9^. Macaques are both terrestrial and arboreal; they are adapted to climbing and leaping through the trees present in their natural habitats^10^, often using vertical routes to escape predators or to find a place for resting^9^. Indeed, wild macaques can spend up to approximately 28% of their awake time moving and travelling and 36% for feeding and foraging^11,12^. These characteristics are a key consideration when designing captive housing to enhance the quality of life for macaques.

Environmental enrichment aims to enhance the welfare of animals by providing opportunities to express natural behavior and to have a degree of choice and control over the environment. Usually, it comes in the form of changes to the structure and content of the housing environment^13^. For macaques, structural enrichment can include the addition of space, ladders, perches, porches, visual barriers, swings, puzzle feeders, or other manipulable objects^14^. These structural modifications promote a more natural and varied behavioral repertoire. Expanding usable vertical space encourages climbing, leaping, and varied locomotion, particularly important in more arboreal species such as the long-tailed macaque^15^. Social management is enhanced through structures that provide cover and opportunities for hiding from aggressive conspecifics. Playing, exploration, and foraging can be stimulated through the use of engaging elements^14^.

While the importance of environmental enrichment is reflected in the legal mandates of animal welfare regulations^16,17^, the actual effectiveness of these interventions in improving the welfare of captive macaques has not been systematically assessed. Although broader meta-analyses have explored various types of enrichment (e.g. social, sensory, or interactive) across different primate species^18,19^, no previous study has focused exclusively on structural enrichment specifically within macaques. This gap in comprehensive, evidence-based evaluation means that despite regulatory pressure to provide enrichment for animals housed in research facilities, there is a lack of clear, consistent data on which specific interventions yield the most significant welfare benefits across different contexts. Therefore, the objective of the present investigation was to conduct an exploratory systematic review and meta-analysis to assess the effect that different structural enrichment interventions, introduced within the home cage, have on the welfare of macaques kept in captivity. Our focus is on the macaque species most used in biomedical research: long-tailed (*Macaca fascicularis*), rhesus (*Macaca mulatta*), stump-tailed (*Macaca arctoides*), and pig-tailed (*Macaca nemestrina*) macaques.

We chose behavior as the primary outcome of interest because it is a readily observable and non-invasive index of welfare^20^. Behavioral changes often serve as the most immediate and practical indicators of an animal’s response to its environment and enrichment interventions. The presence of abnormal behaviors (e.g., stereotypies, self-injurious behavior) or the absence of species-typical behaviors directly signals a compromised welfare state, making them crucial targets for improvement in captive settings^6,13^. Any change in behavior following the implementation of an enrichment intervention can be used as a measure of its positive (or negative) effects. We recognize that welfare is a complex biological state that cannot be fully captured by behavioral measures alone^6,21^. However, including a broader range of welfare indicators was not feasible for this systematic review due to inconsistent measuring and reporting practices across the diverse primary studies. Behavioral data, conversely, was more consistently documented, allowing for a meaningful synthesis of the available evidence on the impact of structural enrichment. Our hypothesis was that structural enrichment interventions would be associated with an increase in the duration or frequency of positive welfare-related behaviors (natural, species-typical behaviors) and a reduction in the duration or frequency of negative welfare-related behaviors (e.g. abnormal behaviors) of macaques kept in captivity. To maintain conciseness here, the specific list of behaviors categorized as positive or negative welfare-related can be found in the methods section of this paper.

## Methods

### Protocol and open access data

The protocol for the current study was prepared *a priori* and registered in Open Science Framework (OSF) under registration DOI https://doi.org/10.17605/OSF.IO/Q8PFS. The report adheres to the PRISMA guidelines for reporting^22^. The supplementary materials contain detailed information regarding the methods used in the study, links to the data repository, and deviations from the registered protocol.

### Search strategy

Four databases were searched for relevant studies: PubMed, Embase (Ovid), Web of Science (core collection and zoological record collection) and the Animal Welfare Institute (AWI) refinement database (https://awionline.org/content/refinement-database). The dates and description of the search are provided in the supplementary material. We were interested in studies that assessed the effect of structural enrichment strategies on the behavior of macaques kept in captivity. For our study, **structural enrichment** was defined as **any structural item and/or space added to the home cage that increases its complexity and/or its usable space**. The reference lists of relevant reviews and of included studies (n = 153) were reviewed as a supplementary method for identifying potentially relevant publications.

### Screening Platforms

The Covidence systematic review software (Veritas Health Innovation, Melbourne, Australia) was used for most of the screening process^23^. The ASReview interface^24^ was employed to streamline the title and abstract screening.

### Screening process

After duplicated publications were removed, the remaining publications were screened in two phases. The initial title and abstract screening was streamlined by employing active learning (using ASReview) ^24^. For a complete account of the process, please refer to the supplementary material. Briefly, an initial random set of 600 references was screened manually by two reviewers to create a dataset to train a machine learning model. One reviewer then screened the remaining references, assisted by the active learning model, until three stopping criteria were met: (1) key pre-selected papers were found; (2) at least 600 records were screened; and (3) 60 consecutive irrelevant records were excluded. To address potential omissions, a deep learning model was used to continue the screening until 60 consecutive irrelevant records were found. Finally, a second reviewer double-screened previously excluded records using a Naïve Bayes model to ensure no relevant studies were missed.

This initial title and abstract screening excluded studies that did not describe an original study in captive macaques that employed a structural enrichment intervention (**Table 1**, criteria a-d). In cases where there was disagreement between reviewers or incomplete information for exclusion, the study was moved on to the next phase. During the full-text screening, two independent reviewers screened the remaining studies. Only studies that fulfilled all criteria specified in **Table 1** were included in our study. These criteria ensured that we only included studies that assessed the effect of structural enrichment strategies on the behavior of macaques kept in captivity. A third reviewer, casting a deciding vote, solved discrepancies during this phase. We categorized behavior (the outcome of interest) into two main types: positive-welfare-related behaviors and negative-welfare-related behaviors. Positive-welfare-related behaviors were defined as those whose increased frequency or duration is generally considered indicative of a positive welfare state. Negative-welfare-related behaviors were defined as those whose increased frequency or duration is generally considered indicative of a negative welfare state. **Table 1** presents the behaviors that were, *a priori*, agreed upon for inclusion in each of these categories.

**Table 1.**
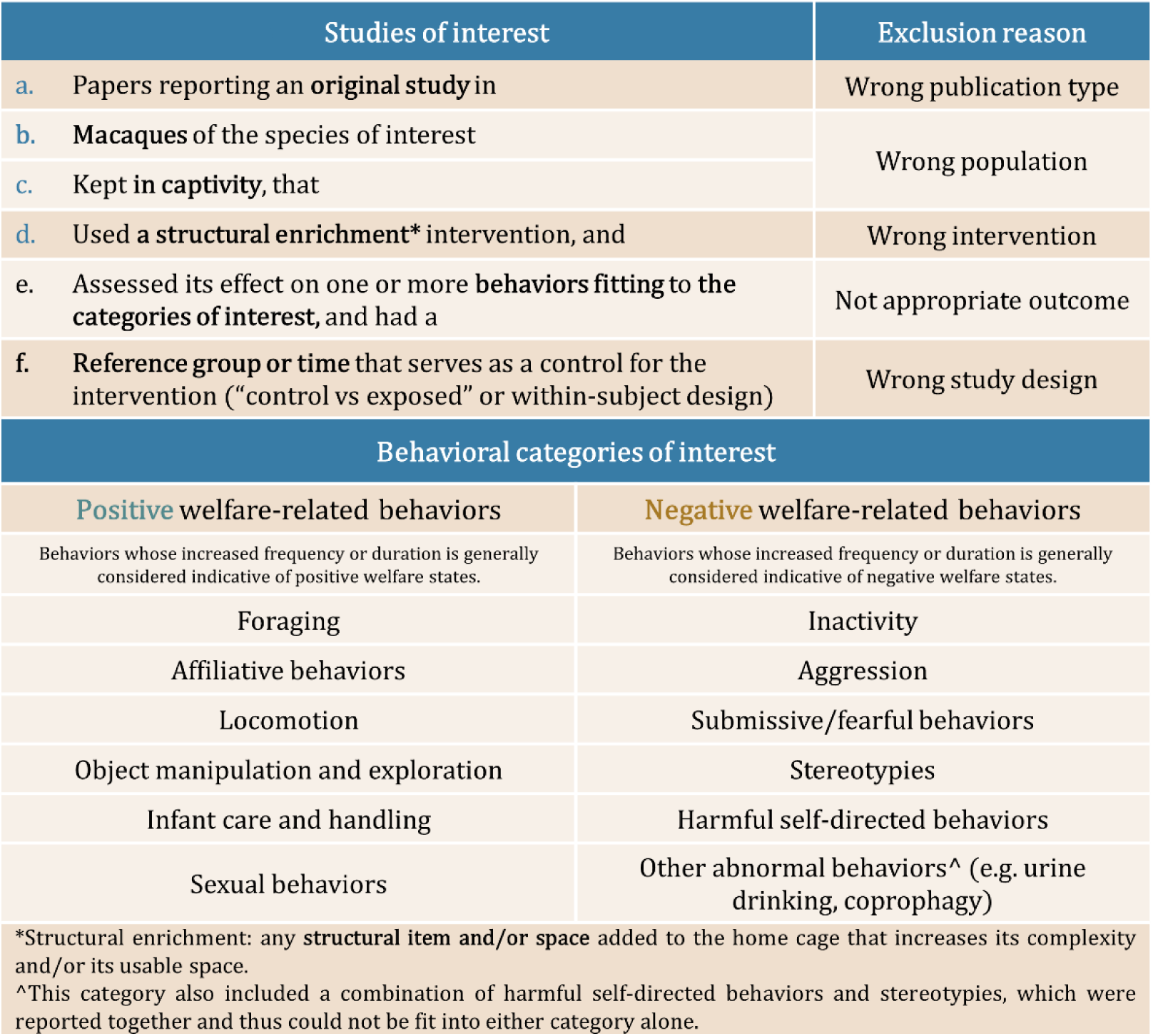
Eligibility criteria and behavioral categories of interest. Criteria *a* to *d* were checked during title and abstract screening, whereas all the criteria were assessed during full-text screening. The reasons for exclusion match those reported in the PRISMA flow diagram. The bottom half of the table details the behaviors that were included in the two types of behavioral categories of interest.

| Studies of interest |  | Exclusion reason |
| --- | --- | --- |
| a. | Papers reporting an **original study** in | Wrong publication type |
| b. | **Macaques** of the species of interest | Wrong population |
| c. | Kept in captivity, that |  |
| d. | Used a **structural enrichment\*** intervention, and | Wrong intervention |
| e. | Assessed its effect on one or more **behaviors fitting to the categories of interest**, and had a | Not appropriate outcome |
| f. | **Reference group or time** that serves as a control for the intervention ("control vs exposed" or within-subject design) | Wrong study design |
| Behavioral categories of interest |  |  |
| Positive welfare-related behaviors |  | Negative welfare-related behaviors |
| Behaviors whose increased frequency or duration is generally considered indicative of positive welfare states. |  | Behaviors whose increased frequency or duration is generally considered indicative of negative welfare states. |
| Foraging |  | Inactivity |
| Affiliative behaviors |  | Aggression |
| Locomotion |  | Submissive/fearful behaviors |
| Object manipulation and exploration |  | Stereotypies |
| Infant care and handling |  | Harmful self-directed behaviors |
| Sexual behaviors |  | Other abnormal behaviors^ (e.g. urine drinking, coprophagy) |
| *Structural enrichment: any **structural item** and/or **space** added to the home cage that increases its complexity and/or its usable space. |  |  |
| ^This category also included a combination of harmful self-directed behaviors and stereotypies, which were reported together and thus could not be fit into either category alone. |  |  |

### Study details and outcome data extraction

**Supplementary table 2** presents the list of study details and outcomes extracted from each included study. If there were more than one experiment in the study that fulfilled the inclusion criteria, each experiment was extracted individually. Thus, each study contributed one or more experiments to our dataset. Both within and between-subjects designed experiments were extracted. Two reviewers independently performed the extraction of the study details, while a third reviewer resolved discrepancies. One reviewer extracted the outcome data (according to previous agreed-upon rules – refer to document hosted on OSF alongside the protocol) while consulting with the rest of the team on the best way to extract the data for more difficult cases. This ensured consistency in the outcome data extraction. We had defined three broad categories of structural enrichment: **Physical enrichment**, which included a cage size increase, or the introduction of furniture or manipulable objects (e.g. perches, visual barriers, porches/verandas/balconies, ladders, barrels); **Feeding (structural) enrichment**, limited to objects incorporated in the home cage that promote foraging (e.g. foraging devices like puzzle balls, foraging boards); and **Mixed enrichment**, incorporating both types of enrichment.

Whenever possible, information was gathered directly from original data, tables, or text. Data presented in figures were obtained using a digitizing tool^25^. Efforts were made to contact authors in case of missing data. If no response was received and other options for obtaining the data were exhausted, the study/experiment was excluded from the meta-analysis.

Given the heterogeneous nature of the research on the topic and flexible designs and reporting, we abstained from performing a structured risk of bias assessment or test for publication bias. We consider our study to be exploratory in nature.

### Data synthesis

#### Meta-analysis

The statistical analysis and visualizations were performed in RStudio 2024.12.1+563^26^ with R version 4.5.0 (2025-04-11 ucrt)^27^. The full list of used packages is presented in the supplementary material. A meta-analysis was considered possible when a minimum of two studies were included per type of enrichment^28^. Whenever the standard error was reported, it was converted to a standard deviation for the meta-analysis. The effect of the enrichment interventions was evaluated by comparing pre and post-enrichment behavior and by comparing behavior in control and exposed animals. Given the nested structure of our data (each study having multiple experiments and each experiment reporting multiple behaviors), we conducted a three-level meta-analysis with robust variance estimation (RVE) for structural enrichment as a whole and for each of the three major types of enrichment (physical, feeding, mixed). The effects of structural enrichment and of the three different types of enrichment were calculated separately for the two major categories of behaviors (calculating one pooled estimate for positive welfare-related behaviors and another for negative welfare-related behaviors). Pooling and weighting of individual effect sizes (standardized mean differences, SMD - Hedges’ g) was done with a random-effects model using the restricted maximum likelihood (REML) for estimating τ^2^ at level 2 (variability between different experiments *within* the same study) and at level 3 (variability between studies). The statistical measure for assessing heterogeneity was I^2^. We conducted sensitivity analyses by re-running the meta-analyses after removing influential studies and by focusing only on studies directly comparing no enrichment to enrichment. Subgroup analyses were conducted to explore whether differences in research facility, the species of macaques, the sex of the animals, housing type, rearing conditions or the subtype of physical enrichment can explain some of the observed heterogeneity in effect sizes across studies. Only the categories that had 2 or more studies were included in these subgroup analyses (**Supplementary table 4)**.

#### Descriptive analysis

Experiments whose complete behavioral data could not be extracted for statistical pooling were included in a descriptive analysis to maximize the utility of all extracted data. For these experiments, and for those included in the meta-analysis, we directly extracted from the full texts whether the behavior of interest increased, decreased, or showed no significant change following the structural intervention. The results were analyzed by type of structural enrichment intervention, by type of behavioral category (**Table 1**), and when only considering those experiments that didn’t have any enrichment during the control/baseline condition.

## Results

### Screening process

The PRISMA flow diagram in **Figure 1** illustrates the steps of our study selection process, including quantity and rationale behind the exclusion of studies at each stage. From 5,785 non-duplicated records found through our search (initial search plus additions found in the AWI database and reference lists), 76 studies described 102 original experiments on captive macaques of the species of interest who were exposed to a structural enrichment intervention. Each experiment reported one or more behaviors, sometimes reporting two or more behaviors belonging to the same category. The complete list of the studies that were included is given in the supplementary material. A total of 427 behavioral outcomes (also referred to as ‘behaviors’) were reported in our 102 experiments.

**Figure 1.**
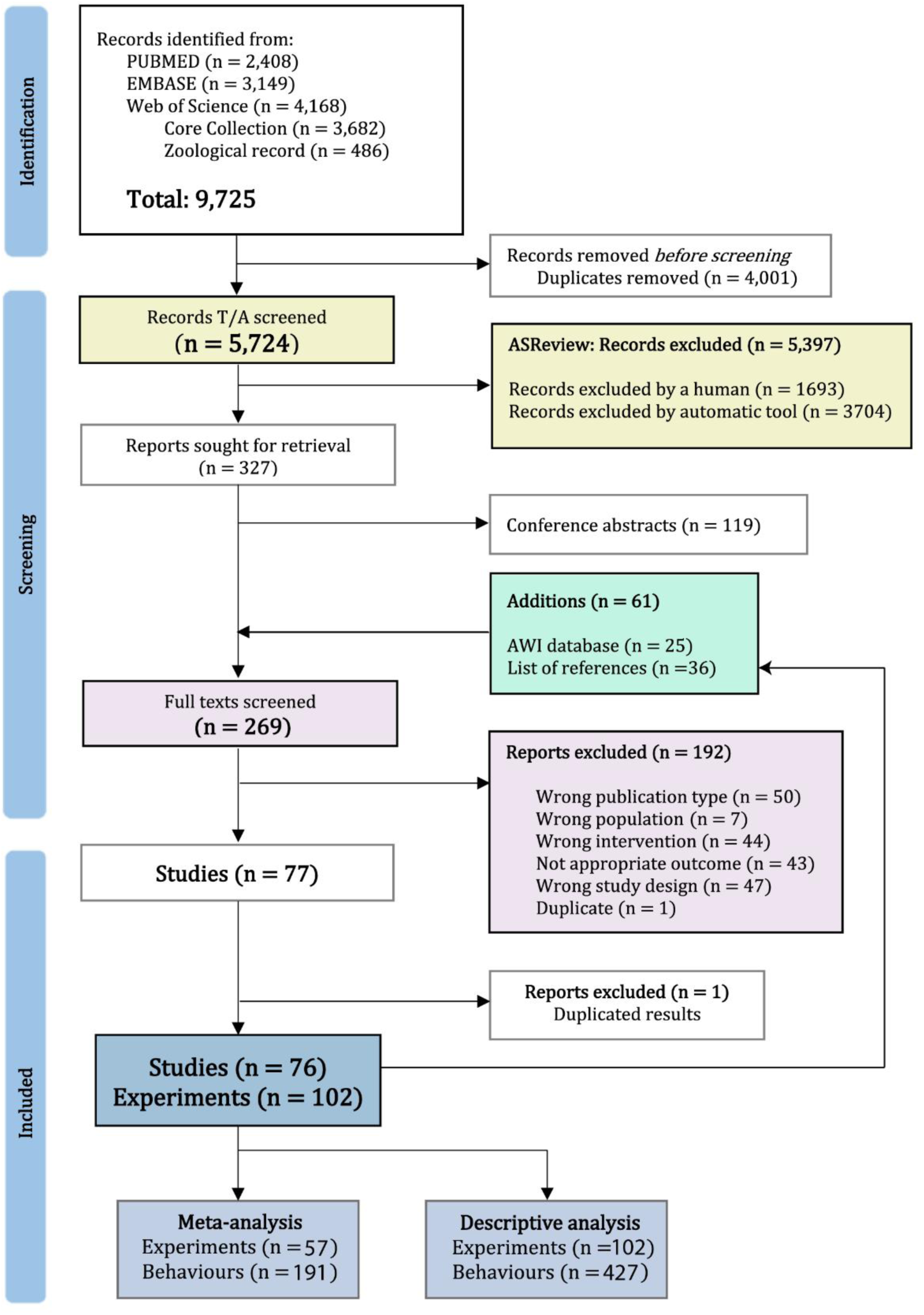
PRISMA flow diagram. Our initial search across databases yielded 9725 reports. Following the removal of duplicates, we screened 5,724 reports by title and abstract using ASReview ^24^. Through this method we screened 35.3% of the records (n = 2020) and included 327 for full-text screening. 61 additional records were identified in the Animal Welfare Institute (AWI) database and through searching in the reference lists of relevant reviews and included studies (n = 153). After excluding conference abstracts, 269 full-texts were screened. Ultimately, 77 papers satisfied all inclusion criteria. We excluded one of these studies because it reported a summary of results already published in previous studies by the same author. In the end, 76 studies, comprising 102 unique experiments, were included. We did not have the complete data to include 45 experiments in the meta-analysis; therefore, their results are included in the descriptive analysis in our study. This diagram is adapted from the PRISMA 2020 guidelines for systematic reviews^22^.

### Characteristics of the included studies

**Figure 2** is an infographic summarizing highlighted characteristics of the studies. The included studies were published between 1963 and 2024. The majority (n = 44) were published between 1990 and 2000 (**Figure S3**). Most of the research originated from the United States (80.3%), carried out in facilities in academic or public research settings (86.8%). 13.1% of the studies originated in countries from Europe. The rest originated in Australia, Canada, Korea and Mexico. Notably, there were no studies from countries in Africa or from other major primate-using and breeding nations in Asia. Only 9.2% of the studies were carried out in a research facility in the private sector (**Figure S4**). Less than 4% occurred in non-research settings (zoos, rescue centers, and sanctuaries).

**Figure 2.**
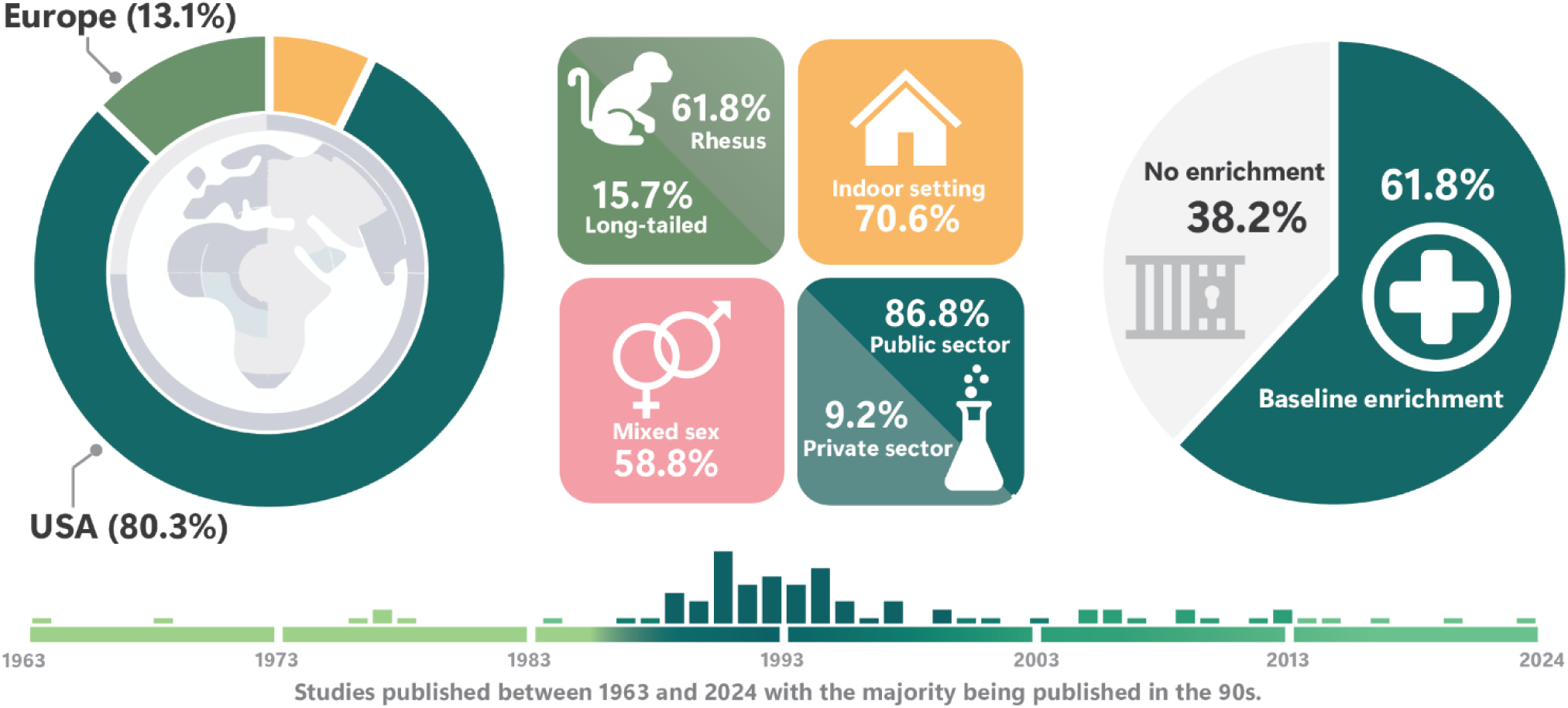
Infographic study characteristics. Summary of key study attributes, including the geographical location of the studies, experimental characteristics and environmental enrichment at baseline (see text for detailed information). The timeline at the bottom shows the distribution of published studies across years (See Figure S3).

#### Subject characteristics

From this point forward, the description will focus on experiments rather than studies.

A large proportion of the experiments (61.8%) employed rhesus macaques, with long-tailed macaques being the second most common species (15.7 %). Pig-tailed macaques and stump-tailed macaques were each used in 9.8% of the experiments (**Figure S4**). A mixed group of different species was used in 2.9% of the experiments. The ages of the experimental subjects varied widely, ranging from infants to elder macaques. In many instances, subjects with different developmental stages were combined in the same experimental group. Over half of the experiments (58.8%) used experimental groups that were mixed in terms of sex, while around a fifth employed groups composed of only males or only females (20.6% and 19.6%, respectively). In 1% of experiments the sex was not reported **(Figure S4)**. Information about the early rearing environment (the first six months of life) was missing for 51% of the experiments. In 20.6% of the experiments, the experimental group had subjects with mixed rearing conditions. In the rest of the experiments, the subjects were either raised in groups (14.7%), raised with their (foster) mother only (2.9%), or in a nursery (3.9%). 6.9% of the experiments studied animals that had been wild-caught. Most of the experiments employed a within-subjects design (n = 76, 74.5%), while only about a quarter (n = 26, 25.5%) used a control versus exposed group design **(Figure S4)**. The subject characteristics for each type of experiment are reported in detail in the supplementary materials.

#### Intervention characteristics

##### Housing

More than half of the experiments studied single-housed macaques (53.9%), followed by group-housed (31.4%) and pair-housed macaques (6.9%). Groups of macaques with varied housing conditions were employed in 7.8% of the experiments. For 70.6% of the experiments, the cages were located indoors, 7.8% were exclusively outdoors, and in 15.7% the captive environment had a portion indoors and a portion outdoors. For most studies, the housing conditions of the animals did not change when the enrichment intervention started, except for six. In two of these studies (4 experiments)^29,30^, the enrichment intervention involved a transition from single, indoor housing to outdoor group settings. In the remaining 4 studies (5 experiments)^31–34^, the transition only involved a shift from an indoor to an outdoor setting. The majority of the cages provided between 0.4 and 0.89 m^2^ floor space per animal. The typical (median) individual space allowance was 0.6 m^2^, but it ranged from a minimum of about 0.13 to a maximum of 26.28 m^2^ per animal. The density of animals varied, with the majority of cages having between 1.1 and 2.5 animals per square meter. The typical density was about 1.6 animals per square meter. It’s important to note that for these calculations, each individual animal was counted equally, regardless of its size or age (e.g., three infants or three adults in the same cage would result in the same density estimates). See supplementary material.

##### Enrichment

Of the 102 experiments included, the majority (61.77%, n = 63) involved some form of baseline structural enrichment upon which the tested enrichment intervention was applied. In contrast, only 38.23% (n = 39) of the experiments had no enrichment in their baseline or control conditions. Notably, all but one of these studies were carried out before 2002, the other being conducted in 2008 in Korea.

**Figure 3** is a chord diagram that shows the number of experiments that employed none or some sort of enrichment at baseline and the type of structural enrichment they tested (e.g. experiments that at baseline had physical enrichment but tested the effect of a mixed intervention). Among experiments with no baseline/control enrichment, 26 investigated the effect of introducing physical enrichment, followed by a mixed intervention (n = 9) and feeding enrichment (n = 4). In 17 experiments, it was not reported whether the macaques had any enrichment to begin with. Most of the experiments that had enrichment at baseline/control, had so in the form of physical enrichment (34.31%, n = 35). Among these studies, the majority tested interventions of adding more physical enrichment (**Figure 3**). The space allowance and density in the enriched conditions had a wider spread than the control conditions. The majority of the enriched cages provided between 0.4 and 2.8 m^2^ per animal. The typical individual space allowance was slightly higher than control conditions (0.7 m^2^), but it ranged substantially more from a minimum of about 0.13 to a maximum of 5,000 m^2^ per animal. The density in the enriched conditions was slightly smaller. The typical density was about 1.4 animals per square meter and most cages had between 0.4 and 2.5 animals per square meter. See supplementary material.

**Figure 3.**
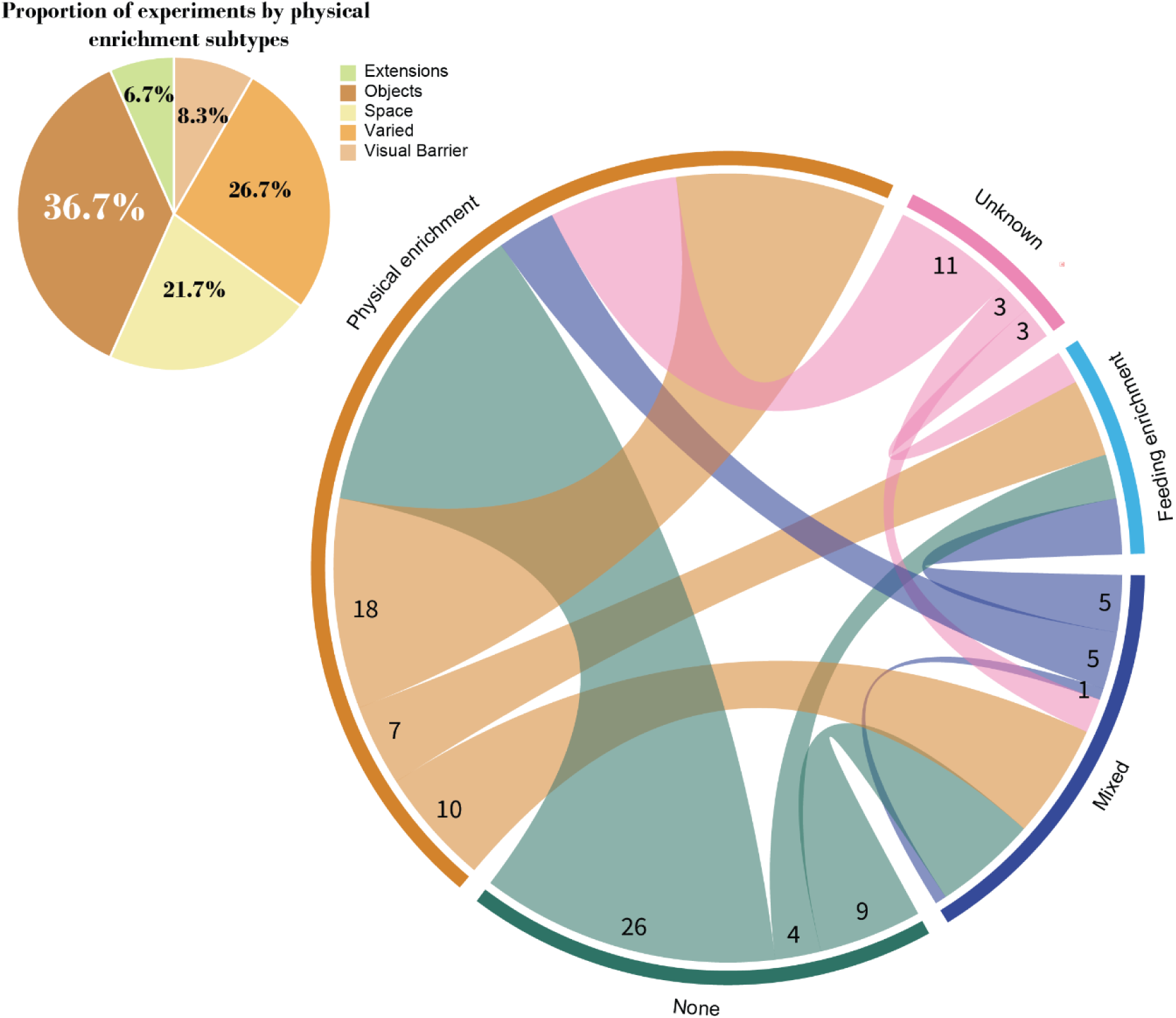
Enrichment at baseline/control and at intervention phases. This chord diagram shows the number of experiments based on the type of enrichment present at baseline/control conditions and the tested type of structural enrichment intervention. The outer ring segments represent the types of structural enrichment (Physical, Feeding, Mixed, None, or Unknown), with the numbers indicating the start of the connection and representing the total count of experiments starting with that condition. The chords connecting these segments end at the intervention types tested in these experiments. For example, among experiments with no baseline/control enrichment (None), 26 investigated the effect of introducing physical enrichment, and 7 of those that had physical enrichment at the start tested the effect of adding feeding (structural) enrichment. The physical enrichment introduced in the intervention phase was further divided into 5 subtypes: Objects, space, extensions, visual barriers, and varied. The pie chart at the upper-left corner shows the proportion of studies that used each of these physical subtypes during the intervention phase (total of 60 experiments).

The category physical enrichment during the intervention phase was further divided into 5 subtypes: *Objects* (e.g. toys, barrels, wood sticks, etc.), *space* (when only an increase in cage size was tested), *extensions* (perches and porches), *visual barriers* (e.g. privacy panels, covers), and *varied* (when more than one of the subtypes above were combined). From the 60 experiments (58.8%) that tested the addition of physical enrichment, 22 added objects to the home cage (36.7%), 13 only increased the size of the cage (21.7%), 4 used extensions, and 5 introduced barriers. The rest used a mix of all these subtypes (n = 16, 26.7%) (**Figure 3**, upper-left corner).

The duration of environmental enrichment across the included studies varied considerably, ranging from a few hours to lifetime exposure. Short enrichment periods (half an hour to six days) were used in 17 experiments. Most experiments (n = 63) tested interventions lasting between one week and seven months, while 8 experiments had durations of 1-11 years. ‘Lifetime’ enrichment was reported in ten experiments, and the duration was unknown in four.

### Data synthesis

#### Meta-analysis

Out of the 76 included studies, 42 studies (57 experiments, 191 behavioral outcomes) had complete data that could be included in a meta-analysis (**Figure 1**). 26 studies reported positive-welfare-related behaviors, while 35 reported negative-welfare-related behaviors. Structural enrichment, combined across types, decreased negative-welfare-related behaviors (number of studies (k) = 35, SMD: -0.32; 95% CI: [- 0.49, -0.15]; *p* = 0.0004) and increased positive-welfare-related ones (k = 26, SMD: 0.31; 95% CI: [0.02,

0.59]; *p* = 0.036). The between-study heterogeneity (I^2^ Level 3) made up an estimated 48.5% of the total variability for positive behaviors, and 16.12% for negative behaviors. The within-study heterogeneity (I^2^ Level 2) could not be estimated (I^2^ = 0%) in either case. The species, sex, housing conditions or the type of research facility (public vs private) did not explain a significant amount of the variation in effect sizes across our studies (**Supplementary table 4**). Rearing condition did not emerge as a significant moderator of effect size variability either, even when the analysis was restricted to abnormal behaviors (harmful self-directed behaviors, stereotypies, and other abnormal behaviors) (**Supplementary Table 4**). However, it is important to note that due to limited reporting on rearing conditions, the sample size for these specific subgroup analyses was substantially reduced. Physical and mixed enrichments significantly decreased negative welfare-related behaviors (SMD: **-**0.24, *p* = 0.027; SMD: **-**0,60, *p* = 0.009, respectively) but did not significantly increase positive-welfare-related behaviors (SMD: 0.17, *p* = 0.22; SMD: 0.09, *p* = 0.67, respectively) (**Figure 4**). The different subtypes of physical enrichment did not differ significantly in their effects and did not explain a statistically significant amount of the variation in effect sizes across our studies (**Supplementary Table 4**). However, mixed physical strategies were associated with a significant reduction in negative-welfare-related behaviors. By contrast, interventions limited to only objects or visual barriers demonstrated a lesser (non-significant) effect on both positive and negative behaviors. Increasing cage space alone (ranging from 0.16 to 106 m^2^ of added space) had no discernible effect on either of these behaviors (**Figure 5**). Feeding enrichment had the lowest count of reported behaviors among the enrichment types. It led to a non-significant increase in foraging behaviors (no other positive welfare-related behaviors were assessed for this type of enrichment) (SMD: 3.6; 95% CI: [-0.76, 7.96]; *p* = 0,08) and a non-significant decrease in negative welfare-related behaviors (SMD: -0.36; 95% CI: [-1.20, 0.48]; *p* = 0.15). The forest plots of the analyses can be found in the supplementary material (**Figure S7**).

**Figure 4.**
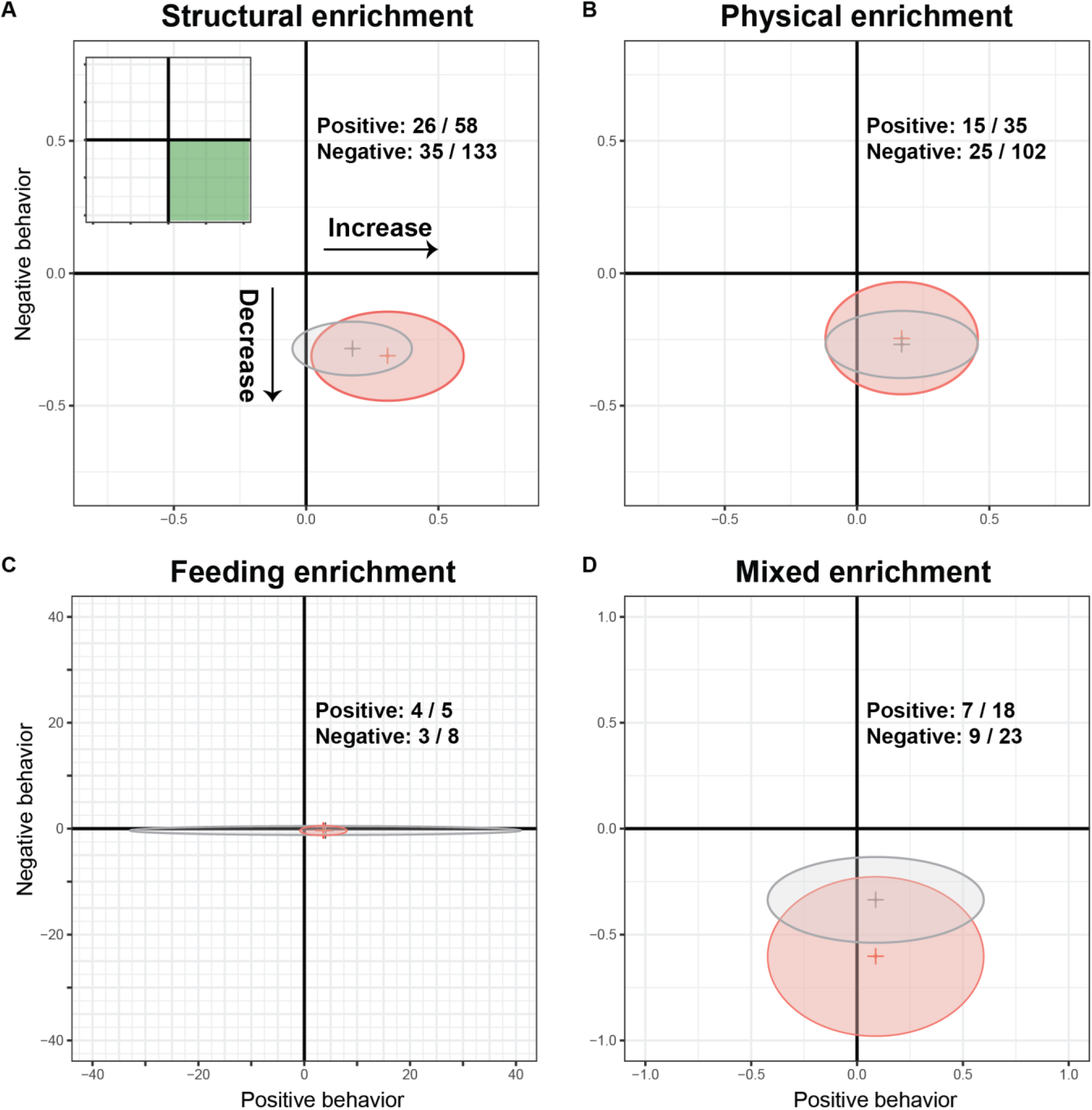
Effect of structural enrichment on behavior. This figure provides a visual representation of the effect of structural enrichment (**A**) and the three different types (**B-D**) on the behavioral domains of interest. The X-axis represents the estimated effect on positive welfare-related behaviors, while the Y-axis represents that on negative welfare-related behaviors. Positive values in either axis indicate an increase in behavior in the enriched condition, while negative values indicate a decrease in comparison to the control group/condition. Interventions that promote positive-welfare-related behaviors and decrease negative-welfare-related behaviors will fall in the lower-right quadrant (**A**, upper left corner), further to the right and down, the bigger the effect. For each type of enrichment, the intersection between the effect estimates on the X and Y-axes is marked with a red cross. The red elliptical regions were created by intersecting the confidence intervals (CIs) of the effect estimates. Regions crossing the zero lines mark a non-significant effect on that specific behavioral domain. The number of studies/behaviors pooled for each intervention are given in the upper right quadrants. The different scatterplots have different scales to accommodate the effect estimates and their CIs. They gray crosses and ellipses represent the shift in effects after removing influential studies (sensitivity analysis).

**Figure 5.**
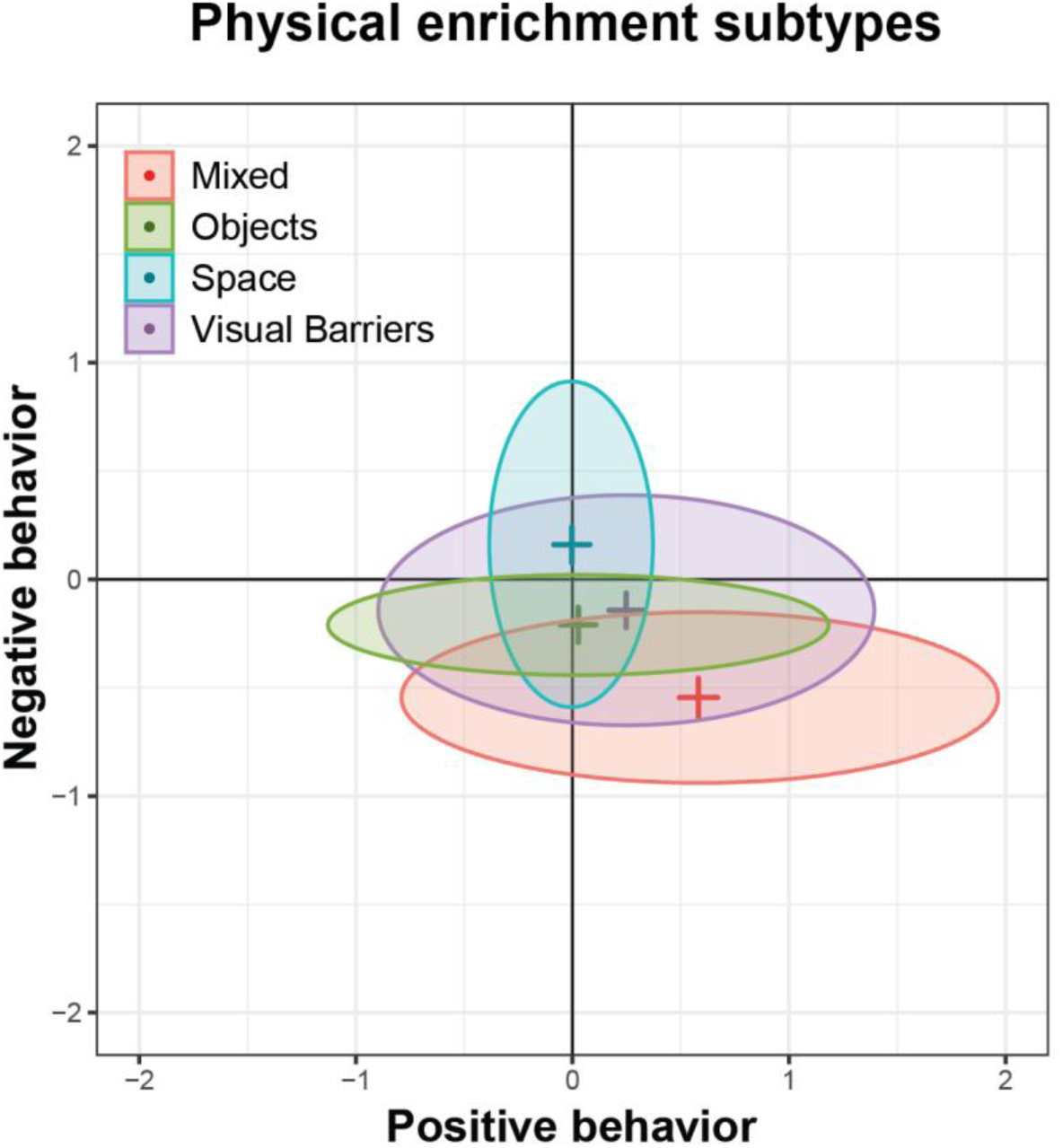
Effect of different physical enrichment subtypes on behavior. This figure provides a visual representation of the effects of different subtypes of physical enrichment on the behavioral domains of interest. The X-axis represents the estimated effect on positive welfare-related behaviors, while the Y-axis represents that on negative welfare-related behaviors. For each subtype of enrichment, the intersection between the effect estimates on the X and Y-axes is marked with a colored cross corresponding to the subtype (legend in the upper left corner). The rectangular regions were created by intersecting the confidence intervals (CIs) of the effect estimates. Increasing space only (light blue) showed a minimal or non-significant positive effect on normal behaviors, while having a non-significant increase in behaviors associated with negative welfare. In contrast, mixing different types of physical enrichment (red) had a noticeable and statistically significant reduction in negative welfare-related behaviors. There was not enough data to run analysis for the subtype extensions.

### Sensitivity analyses

Three positive and three negative behaviors were identified as outliers (**Figure S8**). None represented a data entry error. Two influential studies (contributing 3 experiments) were identified for positive behaviors, both studies contributing results on foraging behavior. Five studies (11 experiments) were identified for negative behaviors, the majority (n = 10) reporting behavioral outcomes related to aggression, and the rest to inactivity, stereotypies and abnormal behaviors (**Figure S9**). Sensitivity analyses were run after removing these influential studies. The effect of structural enrichment, including physical and mixed enrichments, on negative behavior remained significant (**Figure 4**, gray areas), while the effect of structural enrichment on positive-welfare-related behaviors did not (SMD: 0.17; 95% CI: [-0.05, 0.40]; *p* = 0.12).

Our dataset comprised studies that had and did not have enrichment at the baseline. When an enrichment item is *added* to an already enriched environment, its impact might differ compared to its *introduction* in an unenriched setting. Therefore, we re-ran the analyses focusing only on studies that directly compare no enrichment to enrichment. The effect of structural (k = 13, SMD: -0.21; 95% CI: [- 0.30, -0.12]; *p* = 0.0008) and physical enrichments (k = 10, SMD: -0.20; 95% CI: [-0.31, -0.08]; *p* = 0.0062) on negative behavior remained significant; however, the effect of structural enrichment on positive behaviors did not (k = 8, SMD: 0.30; 95% CI: [-0.03, 0.64]; *p* = 0.06). Physical enrichment still led to a non-significant increase in positive welfare-related behaviors (k = 9, SMD: 0.17; 95% CI: [-0.33, 0.67]; *p* = 0.39) (**Figure S10**). There were not enough studies using no enrichment at baseline to perform sensitivity analysis for feeding and mixed enrichments.

#### Descriptive analysis

A total of 427 behaviors were reported in our included studies; 236 behavioral outcomes did not have complete data to be included in our quantitative analysis. However, we did have access to the authors’ interpretation of their results as reported in their papers. These behavioral outcomes together with those included in our meta-analysis were analyzed descriptively. Two thirds of the outcomes were negative welfare-related behaviors (n = 281). Structural enrichment resulted in expected changes in both positive and negative-welfare-related behaviors, though many outcomes remained unaffected. Overall, structural enrichment increased 37.2% of the positive behaviors, while having no effect in 46.2%. Contrary to expectation, 16.6% of positive behaviors decreased following a structural enrichment intervention. Among these positive behaviors, increases were most frequently reported in foraging, locomotion, and object manipulation/exploration, while sexual behaviors showed no increases at all (**Figure 6A**). There was only one report on infant care and handling, and it showed no discernible effect. For affiliative behaviors, structural enrichment had a mixed or inconsistent effect. While there were instances where affiliative behaviors increased, a nearly equal or slightly greater number of instances showed a decrease.

**Figure 6.**
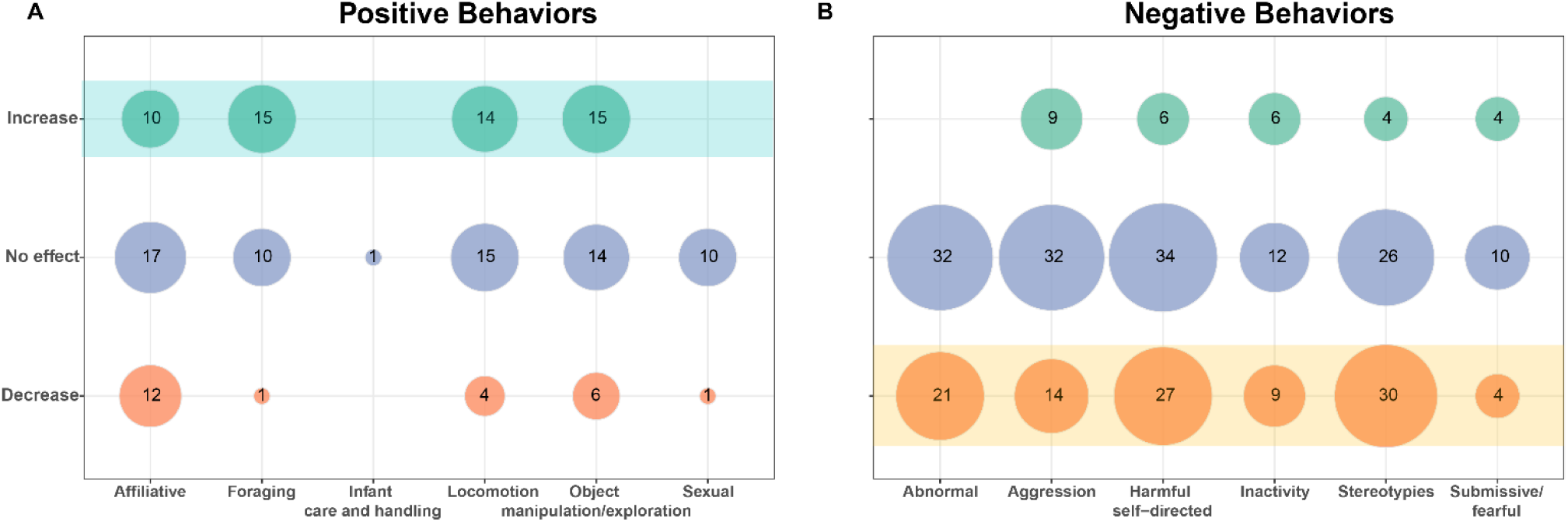
Bubble plots of behavioral changes for structural enrichment. The numbers within the bubbles represent the number of behaviors within each category that were reported to have increased, decreased, or not significantly changed after structural enrichment was introduced into the home cage. Changes in positive-welfare-related behaviors are presented in the left-hand panel, while changes in negative**-**welfare-related behaviors are shown in the right panel. The colored boxes represent the desired effect of enrichment: an increase in species typical behaviors (green) while decreasing those associated with negative welfare (orange).

Structural enrichment decreased 37.5% of negative welfare-related behaviors, while having no effect in 52.1%. A paradoxical increase was observed in 10.4% of these behaviors. Reductions were observed most prominently in stereotypies, harmful self-directed behaviors, other abnormal behaviors, and aggression. However, many negative behaviors belonging to these categories also showed no change (**Figure 6B**). The effect on inactivity and submissive/fearful behaviors is less clear. When only those experiments that had no enrichment during the control/baseline condition were analyzed (47 positive behaviors, 128 negative behaviors), a similar pattern emerged: structural enrichment broadly had either no effect or a positive effect on species-typical behaviors generally considered to be associated with positive welfare, while also having no effect on or decreasing negative-welfare-related behaviors (**Figure S12**).

When the behaviors were split by the type of structural enrichment used (**Figure 7**), we observed that foraging consistently and significantly increased with feeding enrichment. Locomotion showed the most increase with physical enrichment, while object manipulation/exploration was promoted by mixed enrichment. Both physical and mixed enrichments had inconsistent results on affiliative behaviors. Stereotypies, harmful self-directed behaviors and other abnormal behaviors tended to decrease, especially with physical and mixed enrichments. Aggression was reported to decrease mostly with physical enrichment. No specific trends in the effects of subtypes of physical enrichment on positive behaviors were noted (**Figure S13**). However, the presence of mixed physical enrichment items was associated with reductions in harmful self-directed behaviors, stereotypies, and other abnormal behaviors. Individual objects also showed reductions in these categories, though to a lesser degree. Additional space, visual barriers or extensions did not show any clear trends (**Figure S14**).

**Figure 7.**
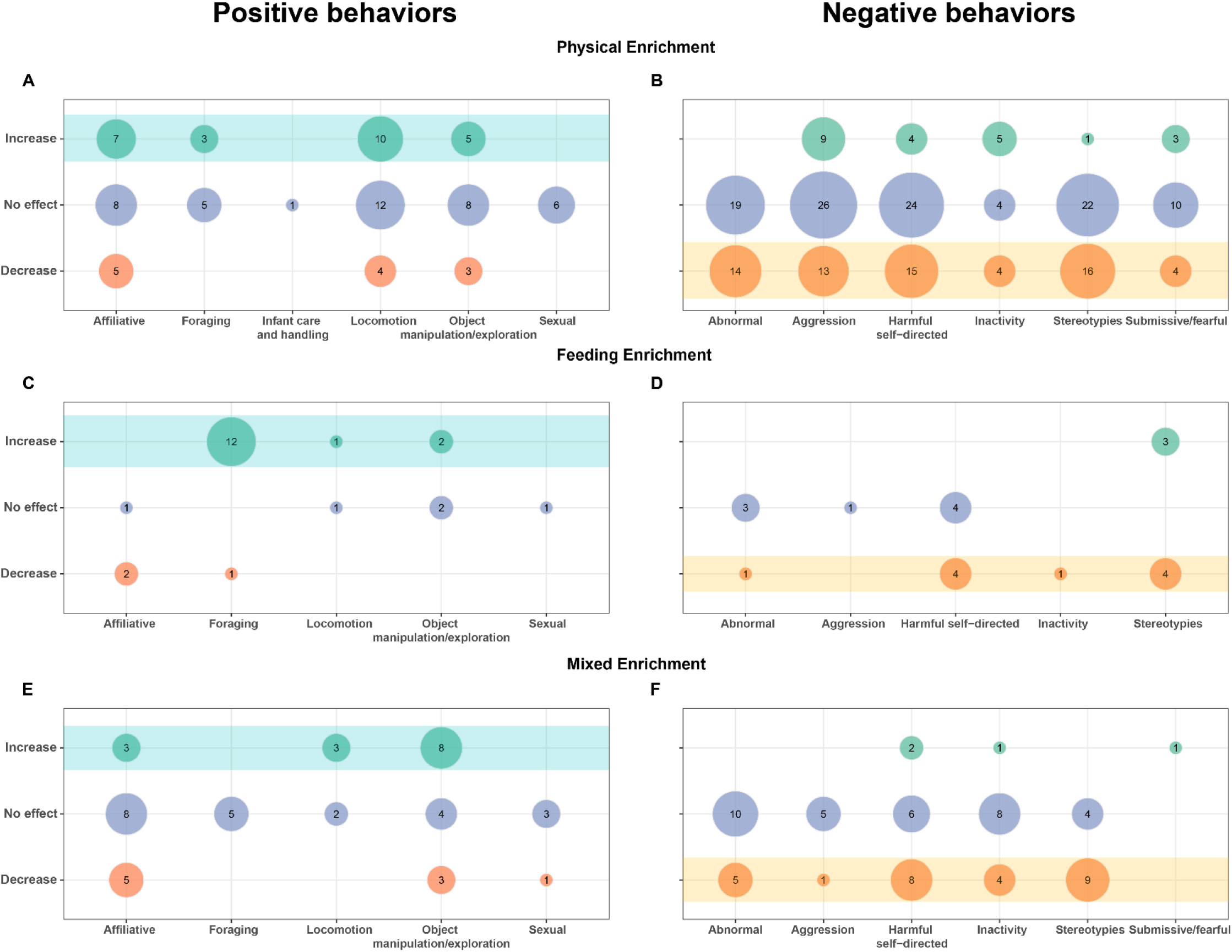
Bubble plots of behavioral changes for different types of structural enrichment. The numbers within the bubbles represent the count of behaviors within each category that were reported to have increased, decreased, or not significantly changed after various types of structural enrichments (physical, feeding, and mixed) were introduced into the home cage. Changes in positive welfare-related behaviors are presented in the left column (A, C, E), while changes in negative welfare-related behaviors are shown in the right column (B, D, F)

## Discussion

This study synthesized 61 years of research on structural enrichment for macaques in research environments, with most publications appearing between 1990 and 2000. The Animal Welfare Act Amendment in 1985 in the United States is arguably a significant driver of this surge in research on the topic. This amendment provided legislative pressure for research facilities to “develop and follow a plan to promote the psychological well-being of primates." Combined with advancements in animal welfare science, the amendment likely increased public demand for ethical treatment of animals. This, along with emerging interests in the effects of enrichment, may have served as a catalyst for research in this area in the late 80s, leading into the 90s^35,36^. Indeed, most of our included studies were conducted in academic or public research facilities in the United States. There were relatively minor contributions from European countries or the private sector. This shift could also explain the disappearance of "no enrichment" control groups after 2002, suggesting that by the early 21st century, the concept of a "barren" control group was no longer considered ethically acceptable.

This regional disparity in the research is not surprising. More restrictive policies on non-human primate (NHP) research in Europe ^37^, coupled with ongoing investment in NHP-dependent disease models and a robust network of National Primate Research Centers in the United States, contribute to a larger U.S. NHP breeding and research capacity ^38,39^. For instance, in 2023, approximately 73,000 NHPs were used for research in the US ^39^, compared to only 7,650 across all European Union member states in 2022^40^. Rhesus macaques, followed by long-tailed macaques, were the species of choice in most studies. This is in alignment with their popularity as model organisms in biomedical researchand use trends in the US^41^. In 2018, the National Institute of Health (NIH) reported that Rhesus macaques comprised 65% of all planned use for project-driven research, followed by long-tailed macaques (15%)^42^.

A substantial portion of experiments exposed the animals to different environmental conditions sequentially, rather than employing independent control and experimental groups. This likely reflects the opportunistic nature of the research: when a veterinarian or welfare researcher implements a housing change, their only option for publishable evidence is often to compare behavior before and after, rarely getting the chance to work with a control group. The characteristics of the study subjects were highly heterogeneous, frequently encompassing wide age ranges and mixing different ages/developmental stages and sexes. Housing conditions also varied considerably. While single housing was most common, nearly half of the experiments incorporated some form of social housing. This heterogeneity in study subjects and housing is expected given the long lifespan of macaques, their frequent re-use in experiments, the regulatory pressure to house them in environments that mimic natural troop structures, and the varying capacities of research facilities to accommodate such social groupings^41^. In fact, animal density in baseline conditions varied considerably in our studies, with a typical density higher than the maximal EU requirements^16^ but still in alignment with US requirements. While some animals in the control conditions were housed in spaces that did not meet US standards, most cages met the minimum requirement of space given to macaques up to 20 kg (Guide for the Care and Use of Laboratory Animals^43^). Some experiments employed very large enclosures that far exceeded minimal requirements. Studies that explored the effect of increasing cage size or shifting housing conditions from indoors to outdoors employed lower animal densities and more individual space during enriched conditions.

### Effect of structural enrichment

Most experiments already included some form of baseline structural enrichment, primarily physical additions rather than more complex enrichment. This means that most studies looked at the effect of *adding* rather than introducing a structural intervention. Physical enrichment (primarily objects) was the most frequently tested intervention, with durations ranging from very short periods to lifetime exposure. Our meta-analysis provides evidence that structural enrichment has a net beneficial effect on the behavior of macaques kept in captivity, thereby strengthening the existing meta-analytical literature that advocates its implementation in primates^18,19^. The beneficial effect was particularly robust in reducing negative-welfare-related behaviors, which were more frequently reported compared to positive-welfare-related behaviors. This focus on negative behaviors likely reflects the historical beginnings of animal welfare science, in which the alleviation of suffering and the prevention of harm were the most pressing issues. Negative behaviors are often clear, observable indicators of distress, frustration, or a poor environment, and therefore carry a strong ethical imperative for immediate intervention^6,13^. This likely directed research towards primarily identifying and reducing these issues at the expense of enhancing positive experiences, even though both are crucial for overall welfare.

Our analysis demonstrated that structural enrichment consistently decreased negative behaviors even after excluding potentially biasing (influential) studies, when the analysis was limited to studies that *introduced* enrichment, or when the analysis included studies that *added* further enrichment to an already enriched environment. This is a crucial finding, as it suggests that the benefits of enrichment are not limited to initial improvements from deprived conditions, but can continue to enhance welfare even in settings that already meet some enrichment standards. Reductions in negative behaviors were most prominent in abnormal behaviors, which included stereotypies and harmful self-directed behaviors, and aggression. This pattern may very well indicate that enrichment is particularly effective in reducing these types of behaviors; however, it is also plausible that research has been more heavily focused on settings where these behaviors were already a significant concern. Thus, it is worth keeping in mind that this pattern may reflect a bias of context for studies rather than a universal effect of structural enrichment. When we looked at different types of enrichment, only physical (mixed physical strategies in particular) and mixed enrichments appear to be effective in mitigating negative-welfare-related behaviors. Feeding enrichment or single physical enrichments did not demonstrate a significant effect. These findings reinforce the prevailing view that enrichment is useful in reducing abnormal behavior^18,44,45^. Considering enrichment is believed to work by allowing animals to engage in preferred activities (which can displace abnormal behaviors), by providing sensory stimulation, and/or by giving them greater control of their environment (like the ability to hide)^13,18^, it follows that more complex or varied enrichment would be more effective. Such complex enrichment would likely be better at reducing underlying frustrations, mitigating contributory stress levels, and/or simply occupying time with preferred behaviors through increased diversity of physical structures^18^.

Indeed, we observed that simply increasing (unfurnished) space provided no discernible benefit. This highlights that meaningful enrichment extends beyond mere spatial expansion, requiring the provision of complex and engaging elements ^46,47^. To truly allow the animals to use such space, it is essential to furnish it with structures that make it both accessible and safe. For example, introducing elements that allow access to the vertical dimension of the enclosure or that can provide crucial refuge from dominant cage mates. In this way, complex environments also tend to reduce stress by providing more options for increased control over the environment^18^. The animals may not take advantage of all the opportunities provided, but those choices still exist. This enhanced control, particularly the ability to influence one’s surroundings and choose interactions, is argued to be a key component of effective enrichment^48^. However, it’s worth noting that enrichment doesn’t always uniformly affect all abnormal behaviors^45^, a pattern also observed in our data. This differential effectiveness suggests that various types of environmental deficits may be responsible for distinct forms of abnormal behavior, implying that more diverse or complex enrichment strategies would be more effective in addressing the full spectrum of these issues^18^. Thus, while the "shotgun" approach of simultaneously presenting multiple enrichment types can complicate the interpretation of study findings^19^, as it makes it difficult to tease apart the individual effects of different enrichment types and thoroughly understand what truly works or does not, its potential for a net benefit to animal welfare still makes it a valuable strategy. This is particularly important in a biomedical context, where ensuring the well-being of healthy animals can also contribute to obtaining reliable and high-quality data^6,49^.

Even though the historical focus has been on mitigating negative states, there is a growing shift towards promoting positive animal welfare states^21^. Our analysis reflects this historical focus, with positive welfare-related behaviors accounting for only a third of our total data. This smaller subset of studies might be one reason why the effects of promoting a normal behavioral repertoire were not as robust as those observed for abolishing negative behaviors. However, it is crucial to remember that decreasing negative behaviors is not the only goal of enrichment. Instead, the ultimate goal is to create environments that allow animals to express a full range of their natural behaviors. Our findings, though less consistent in this area, suggest that enrichment can also help foster a broader, more natural behavioral repertoire. For a primate in captivity, a small positive change can translate into significant improvements in lifetime well-being ^49^. For instance, a small percentage increase in locomotion could mean an animal spends many more minutes per day engaged in natural movement, which is crucial for physical and mental health.

### Varied response to structural enrichment

In our meta-analysis, we observed that a substantial portion of the variability in effect sizes stemmed from the inherent imprecision of the primary studies themselves. However, a significant amount of the variance was due to true differences in the effects across studies: 48.5% for positive-welfare-related behaviors and 16.1% for negative behaviors. Unfortunately, we found no consistent evidence that key characteristics (e.g., age, sex, species, etc.) explained these differences. This does not necessarily mean these factors had no influence; rather, the observed variation was not large enough to account for a significant portion of the overall heterogeneity in effect sizes. It is also possible that for some subgroups we analyzed, we simply lacked sufficient statistical power due to limited data^50^. For example, there was a disproportionately higher number of studies focusing on a specific species, and most studies used mixed-sex groups. This imbalance in sample size across subgroups might have obscured the true impact of the analyzed variables. A critical limitation in our studies was the significant lack of reporting on early rearing environments, given that parental neglect and early life adversity are known to contribute to the development of maladaptive behaviors, which can be resistant to interventions like enrichment^6,45,51^. With our very limited data, we could not find evidence that different rearing conditions explained differences in the effect of enrichment on abnormal behaviors. Similarly, despite other studies suggesting that social conditions can influence enrichment outcomes^19^, we were unable to detect such effects in our analysis.

Another important factor that likely contributed to the unexplained heterogeneity in our statistical model was the duration of enrichment exposure across studies. Some animals experienced lifetime enrichment while others spent only a few hours in the enriched environment. Given that the effect of enrichment can be highly variable depending on its duration^19,49^, this variability is a significant consideration. Our descriptive analysis confirmed heterogeneity in effects across different behaviors. While overall trends were positive, many individual behaviors remained unaffected. This is to be expected. We would not anticipate every behavior in an animal’s repertoire to be influenced by a specific enrichment. For instance, a study might measure locomotion even if the provided enrichment (for example a food puzzle) was primarily designed to affect foraging behaviors. A lack of change in one behavior does not indicate a failed study or ineffective enrichment; it may simply reflect the specific focus of the enrichment and the breadth of the ethogram used for observation.

Structural enrichment most frequently increased foraging, locomotion, and object manipulation/exploration, behaviors that are often considered indicators of engagement and activity. Reductions in negative behaviors were most prominently observed in abnormal behaviors such as stereotypies and harmful self-directed behaviors, and aggression. Importantly, in a small percentage of cases (around 10-16%), enrichment had unexpected (undesirable) effects. These "unexpected" results - the lack of change as well as opposite effects - can have several explanations. One key factor is that behavioral changes are often specific to the type of enrichment provided. For instance, foraging enrichment specifically enhanced foraging behaviors, and we would not necessarily expect it to boost affiliative behavior. In fact, we observed that its impact on other behaviors was less clear. Nevertheless, the absence of an observed increase in other behaviors does not negate the enrichment’s effectiveness within its intended context. Similarly, the absence of a reduction in negative behaviors does not mean the enrichment lacked other welfare effects, as many studies did not measure all relevant behaviors (e.g., only focused on abnormal behaviors). Another explanation is that decreases in positive behaviors may have resulted from shifts in the overall time budget in the presence of enrichment: an animal that was engaged in a lot of affiliative behavior pre-enrichment may have spent more time in enrichment-related activities instead of interacting with peers. This could directly explain why enrichment showed inconsistent effects on affiliative behaviors in our data. Additionally, there is the subject of individual responses, where an animal’s unique personality, housing conditions, past experiences, and current needs can significantly influence how it interacts with and benefits from enrichment ^52,53^. Indeed, long-tailed macaques have shown a greater stress response to laboratory settings than other macaque species used in research^47^. Some animals may potentially find novelty or unpredictability frightening or stressful, or some enrichments might inadvertently cause resource-defense aggression^13,54^.

The array of possible sources of variation in effects underscores the inherent complexity of behavioral responses to environmental interventions. This complexity highlights why a "one-size-fits-all" approach to enrichment is unlikely to be optimally effective. Therefore, those implementing enrichment strategies should be prepared to persevere and try multiple approaches, carefully monitoring individual animals to assess the effectiveness and suitability of each intervention^13^. Ultimately, understanding and accounting for individual differences and the nuanced interplay of various factors will be crucial for maximizing the welfare benefits of structural enrichment.

#### Conclusions

This systematic review and meta-analysis, synthesizing 61 years of research highlights the critical role of structural enrichment in improving the behavioral repertoire of captive macaques. Our findings provide strong evidence that structural enrichment has a net beneficial effect on welfare as measured through behavior, particularly in reducing abnormal behaviors and aggression. Specifically, mixed enrichments that incorporate varied physical enrichment and feeding strategies proved most effective in mitigating these issues. Simply increasing available space, as opposed to usable space, did not have any discernible benefit. While significant progress in primate enrichment strategies has been made, more research is needed on enrichment strategies that consistently promote positive welfare behaviors. Crucially, interpreting the efficacy of structural enrichment requires careful consideration of its context, including factors such as the animals’ individual backgrounds, housing conditions, and the specific duration and type of enrichment provided. Our findings underscore the importance of moving beyond simplistic approaches to enrichment, instead embracing varied strategies that consider the complex behavioral needs of these intelligent animals.

## Author’s Contributions

Jenny Berrio and Otto Kalliokoski conceived and planned the systematic review with help from Dana and Sarah. All authors carried out the review, discussed the results and contributed to the final manuscript. Jenny Berrio performed the statistical analyses and created the data visualizations.

