## Supplementary material for "Structural enrichment for captive macaques - a systematic review and meta-analysis of behavioral outcomes"

Jenny Paola Berrio Sanchez, M.D., Ph.D.; Christopher Musembi Makau, Ph.D.; Dana Pfefferle, Dr. rer. nat.; Sarah Neal, Ph.D.; Otto Kalliokoski, Ph.D.

#### Methods

##### Registration and open access data

Access to the code, additional files and data sets can be found here: https://osf.io/4jemq/

Throughout the text you will find references to specific files found in this OSF repository.

#### Deviations from the pre-registered protocol

- Due to the extensive number of citations retrieved from our systematic search, we opted to introduce a text mining strategy designed to substantially diminish the significant workload associated with manually double screening such a large number of references[1]. Details are given below under "AI-assisted title and abstract screening".
- Rearing, length of the enrichment intervention, and the distinction between research facilities in academia/public institutions and those in companies/private institutions were added to the list of study details extracted from the included studies.
- To address time constraints, the process was changed so that one reviewer, instead of two, extracted the behavioral outcomes for each experiment while still consulting the reviewing team on the best way to extract the data for difficult cases. This was viewed as a good compromise preventing an unnecessarily lengthy process while still aiming for consistency in data extraction.
- Subgroup analyses, initially planned for each specific type of enrichment, were instead conducted using the entire dataset. This decision was driven by the very limited number of studies available for the 'mixed' and 'feeding' enrichment categories, which precluded reliable independent analysis of these subgroups. Using the full dataset for the subgroup analysis allowed greater statistical power and more robust overall results. Additional subgroup analyses by sex, rearing condition and subtype of physical enrichment were performed to complement our exploration of heterogeneity in our data.

- Additional sensitivity analyses were run after removing data that were modified to calculate effect sizes and after identifying influential studies (see “Modified data” and “Outliers and influential cases in the Meta-analysis section).
- Instead of doing a descriptive analysis of only those experiments whose results could not be included in the meta-analysis, we decided to include all the behaviors extracted in the descriptive report (see “Descriptive analysis” section). This option was preferred because it maximized the utility of all extracted data, ensuring that every piece of behavioral information contributed to the understanding of the literature, even if it could not be statistically pooled. Additionally, this approach prevents the quantitative and qualitative analyses from conflicting.

#### Search strategy

For Pubmed, Embase, and Web of science, studies were retrieved on the same day (July the 15, 2024) without any imposed constraints. Each search string was composed of keywords and indexed terms, which were tested and improved to eliminate terms that yielded no results or significantly decreased the accuracy of the search (see Refining the search terms.docx). The final two strings (“a” and “b” below) were merged to obtain the list of potential studies for each database. The first string (a) aimed to identify studies concerning the macaque species of interest. The second string (b) comprised search terms pertaining to welfare, housing, and enrichment.

The complete searches were as follows:

##### *Pubmed*

(a) ("Macaca"[Mesh] OR "macaca*"[tiab] OR "macaque*"[tw] OR "Macaca fascicularis"[Mesh] OR "cynomolgus"[tw] OR "crab-eating"[tiab] OR "long-tailed"[tiab] OR "Macaca mulatta"[Mesh] OR "rhesus"[tw] OR "Macaca arctoides"[Mesh] OR "stump-tailed"[tw] OR "bear macaque"[tiab] OR "Macaca nemestrina"[Mesh] OR "pig-tailed"[tw] OR "pigtail"[tiab]).

(b) ("Animal Welfare"[Mesh] OR "welfare"[tw] OR "well-being"[tw] OR "Housing, Animal"[Mesh] OR "housing"[tiab] OR "enclosure*"[tiab] OR "cage*"[tiab] OR "caging"[tiab] OR "enrichment"[tw] OR "enriched"[tw] OR “porch”[tiab] OR “porches”[tiab] OR “perch”[tiab] OR “perches”[tiab] OR “veranda*”[tiab] OR “balcon*”[tiab] OR “visual barrier*”[tiab] OR “privacy panel*”[tiab] OR “puzzle*”[tiab])

##### *Embase*

(a) macaca.ti,ab. or exp Macaca arctoides/ or exp Macaca/ or exp Macaca fascicularis/ or exp Macaca nemestrina/ or macaque*.mp. or cynomolgus.mp. or crab-eating.ti,ab. or long-tailed.ti,ab. or macaca mulatta.mp. or exp rhesus monkey/ or rhesus.mp. or stump-tailed.mp. or bear macaque.ti,ab. or pig-tailed.mp. or pigtail.ti,ab.

(b) exp animal welfare/ or exp welfare/ or exp experimental animal welfare/ or exp wellbeing/ or well-being.mp. or exp animal housing/ or housing.ti,ab. or enclosure*.ti,ab. or exp cage/ or cage*.ti,ab. or caging.ti,ab. or exp environmental enrichment/ or enrichment.mp. or enriched.mp. or porch.ti,ab. or porches.ti,ab. or perch.ti,ab. or perches.ti,ab. or veranda.ti,ab. or balcon*.ti,ab. or visual barrier.ti,ab. or privacy panel.ti,ab. or puzzle*.ti,ab.

##### *Web of science*

a) TS=("Macaca" OR "macaca*" OR "macaque*" OR "Macaca fascicularis" OR "cynomolgus" OR "crab-eating" OR "long-tailed" OR "Macaca mulatta" OR "rhesus" OR "Macaca arctoides" OR "stump-tailed" OR "bear macaque" OR "Macaca nemestrina" OR "pig-tailed" OR "pigtail")

b) TS=("Animal Welfare" OR "welfare" OR "well-being" OR "Housing, Animal" OR "housing" OR "enclosure*" OR "cage*" OR "caging" OR "enrichment" OR "enriched" OR "porch" OR "porches" OR "perch" OR "perches" OR "veranda*" OR "balcon*" OR "visual barrier*" OR "privacy panel*" OR "puzzle*")

The AWI refinement database was searched on September 17-19, 2024. This database was searched in two phases. First, it was filtered using the terms ‘Macaque’, ‘Environmental enrichment’, ‘Housing’, and 'Welfare assessment'. Subsequently, it was searched using the keywords 'macaque enrichment' due to a website notice indicating that filtering was only reliable for publications after 2010 at the time of the search. A total of 25 additional relevant studies were found and full-text screened.

Publications identified in the AWI database and found through reference lists underwent full-text screening.

#### AI-assisted title and abstract screening

Active learning is an iterative machine learning approach where an algorithm interactively learns to identify relevant records by engaging with a reviewer (human-in-the-loop)[1]. An initial labeled dataset (prior knowledge) primes the model, which then iteratively queries the researcher on records it deems likely to be relevant. The researcher's input retrains the model, a process that continues until a set of stopping criteria are met.

We used the "ASReview" software[2] for the active learning-based screening given its open source code and ability to compare training models. We conducted this screening phase throughout August 2024. The stopping criteria was determined in accordance with the SAFE procedure[3]:

1. **Phase 1: Initial Random Screening.** To establish the **prior knowledge**, we performed a manual dual screening on a random sample of 600 references (10.48% of the total). This screening included 31 references that were retained for full-text screening. Based on this number, we estimated the Fraction of Relevant Records ($F_{RR}$) at **5.2%.** This gave us an estimated number of relevant records ($N_{RR}$) of 296.

2. **Phase 2: Screening via active learning.**
   2.1. *Selection of the active learning model*: To identify the optimal model for our dataset, we employed the simulation module within ASReview. The 'prior knowledge' dataset was used in two sets of simulations. Initially, we compared the performance of various training models on our data, evaluating them based on three metrics[4]:

   - **Work Saved over Sampling** (WSS): This metric quantifies the reduction in screening workload at the cost of failing to detect 5% of the relevant records (recall of 95%). It represents the proportion of irrelevant references correctly excluded by the model.
   - **Extra Relevant Records Found** (ERF 0.1): This metric measures the additional number of relevant records identified by the active learning model compared to random sampling after screening 10% of the dataset.
   - **Average Time to Discovery (**ATD**):** This metric represents the mean number of records screened before a relevant paper is identified, averaged across all relevant records.

   Higher WWS and ERF 0.1, and lower ATD, indicates a better performing model. From our simulations, two models were selected based on their performance metrics (**Figure S1**).

   In a subsequent series of simulations, we further evaluated the performance of the selected models by varying the initial "prior knowledge." Specifically, we conducted 31 simulations, where each simulation utilized a different single relevant study as the initial positive example, while the set of initial irrelevant records remained consistent across all simulations. This allowed us to assess the robustness of the models to different starting points of relevant information (**Figure S2**).

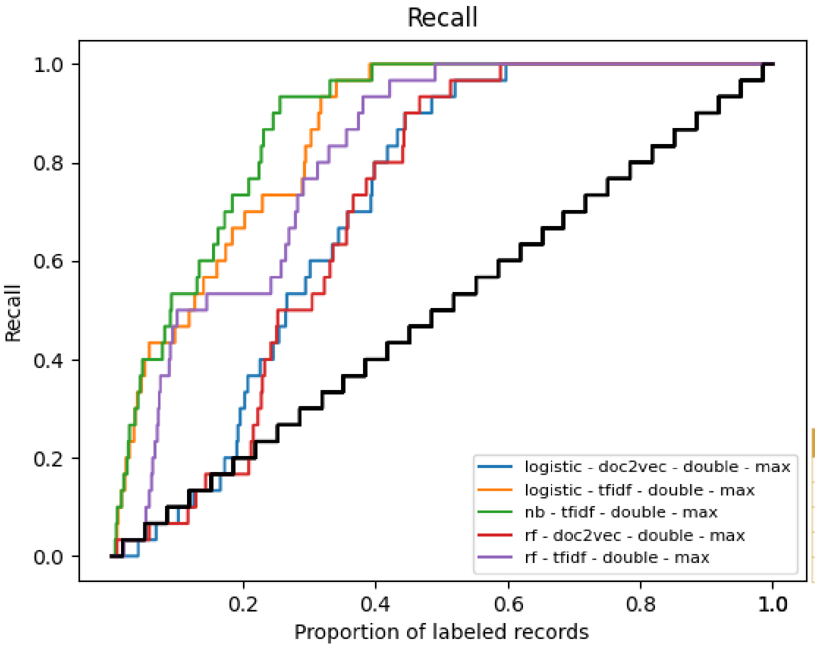


| WSS | ERF 0.1 | ATD |
|---|---|---|
| 0.43 | -0.03 | 172.13 |
| 0.60 | 0.37 | 87.03 |
| 0.66 | 0.43 | 74.03 |
| 0.45 | -0.03 | 175.6 |
| 0.54 | 0.37 | 114.37 |

**Figure S1. Recall performance of various active learning models for our prior knowledge dataset.** The x-axis depicts the cumulative proportion of records reviewed and labeled, while the y-axis shows the corresponding recall achieved by each model (colored lines) in comparison to random screening (black line). Six distinct models were compared: logistic regression with doc2vec embeddings, logistic regression with TF-IDF features, Naive Bayes with TF-IDF features, Random Forest with doc2vec embeddings, and Random Forest with TF-IDF features. The table embedded on the right side provides the performance metrics for each model. ***Naive Bayes with TF-IDF features*** (green line) ***and logistic regression with TF-IDF features*** (orange line) performed better than other models.

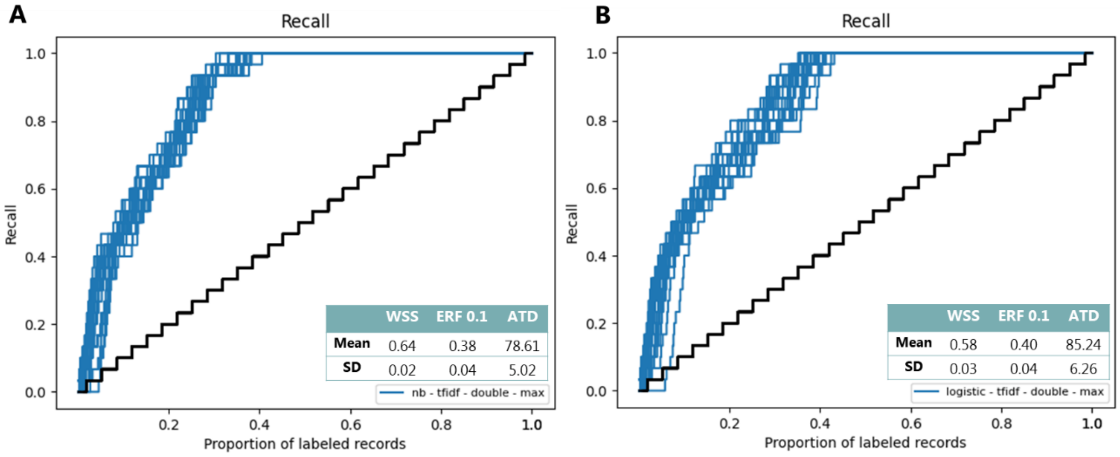


**Figure S2. Recall performance varying prior knowledge.** The x-axis represents the proportion of labeled records, and the y-axis represents recall. Each line in subfigures A and B represents the performance of the models across 31 simulations, each with a different relevant record used as prior knowledge. (**A**) Shows the performance of Naive Bayes with TF-IDF features. (**B**) Shows the performance of Logistic regression with TF-IDF features. The tables embedded in the figures provide the mean and standard deviation (SD) across the 31 simulations for each metric of interest. The model with the best performance for our data was the Naïve Bayes classifier model.

2.2. *Screening*: Using the model with the best performance and using the 600 references manually screened as prior knowledge, one reviewer screened the reminder references until the following **stopping criteria** were met:

a. All the following key papers (used to validate the search strategy) were included:

- Clarence, W. M., et al. (2006). "Use of enclosures with functional vertical space by captive rhesus monkeys (Macaca mulatta) involved in biomedical research." J Am Assoc Lab Anim Sci 45(5): 31-34.
- Paulk, H. H., et al. (1977). "Abnormal behavior in relation to cage size in Rhesus monkeys." J Abnorm Psychol 86(1): 87-92.
- Reinhardt, V. (1990). "Time budget of caged rhesus monkeys exposed to a companion, a PVC perch, and a piece of wood for an extended time." Am J Primatol 20(1): 51-56.
- Reinhardt, V. (1993). "Promoting increased foraging behavior in caged stump-tailed macaques." Folia Primatol (Basel) 61(1): 47-51.
- Gottlieb, D. H., et al. (2015). "Evaluation of environmental and intrinsic factors that contribute to stereotypic behavior in captive rhesus macaques (Macaca mulatta)." Appl Anim Behav Sci 171: 184-191.
- Griffis, C. M., et al. (2013). "Play caging benefits the behavior of singly housed laboratory rhesus macaques (Macaca mulatta)." J Am Assoc Lab Anim Sci 52(5): 534-540.

b. At least 600 records were screened with this method.
c. No relevant records were identified in 60 consecutive screenings.

We found all the key papers after screening 587 papers. We stopped this phase after screening 1261 references.

3. **Phase 3: Switching the model.**
To mitigate the risk of potential omissions of relevant references due to textual ambiguity, we transitioned to a deep learning model. This model employed doc2vec for feature extraction and a fully connected neural network (with two hidden layers) as the classifier. Screening continued until no new relevant records were identified in 60 consecutive screenings. At the end of this phase, we had manually screened 1921 references.

4. **Phase 4: Double screening and quality check.**
To verify that no relevant references were erroneously excluded, a second reviewer re-screened the references initially classified as irrelevant. This phase employed the Naïve Bayes classifier model, and the prior knowledge consisted of the ten highest-ranked and ten lowest-ranked references from the initial screening phase. This process continued until 60 consecutive screenings yielded no additional relevant records. The second reviewer included 3 additional references and after 60 consecutive irrelevant records, he stopped.

In the end, 2020 references were screened employing this method (**35.29%** of the total); 327 were included for full-text screening and 1693 excluded. Active learning reduced our title and abstract screening process by 64,7%.

### Eligibility criteria

The full-text of the studies that passed the title and abstract screening were assessed against pre-defined criteria for inclusion and exclusion (see Protocol.doc). **Supplementary table 1** presents the reasons for exclusion during screening.

### Reference list check

The reference lists of relevant reviews found through the screening process and of the final included studies were reviewed for studies that might have been potentially missed with the initial search. The title of each reference was scrutinized and if it was of potential interest, the paper's abstract and full-text were reviewed. If the study was found to be relevant, the article was full-text screened in the same way as those papers identified through the database search. 36 relevant studies were identified this way.

**Supplementary table 1**. Reasons for exclusion during screening

| | **EXCLUDE** |
|---|---|
| **Not** an **original study** | Literature reviews, editorials, commentaries, opinion pieces, conference proceedings |
| Study does **not** use **live macaques** | Clinical studies, systematic reviews, or in vitro/in silico studies, studies employing animals different from macaques |
| **Species** of macaques **not of interest** | Macaca munzala, Macaca assamensis, Macaca Sylvanus, Macaca radiate, Macaca nigra, Macaca cyclopis, Macaca nigrescens, Macaca hecki, Macaca fuscata, Macaca Silenus, Macaca maura, Macaca leonine, Macaca pagensis, Macaca siberu, Macaca thibetana, Macaca tonkeana, Macaca sinica, Macaca leucogenys |
| Macaques **not** in **captivity** | Macaques living in the wild or in wild-like settings (e.g. safari parks, or wild parks, sanctuaries set up as safari parks) |
| **Other** types of **enrichment**:<br><br>If the study doesn't use any structural enrichment or uses it but doesn't report its individual effect. | • Social<br>• Sensory enrichment (e.g. television or other visual displays, slides, mirrors outside the cage, music or other sounds, odors)<br>• Foraging enrichment (**varied food items or foraging substrates**)<br>• Cognitive (computer-based cognitive tasks including the use of tablets)<br>• Positive reinforcement training or any other type of positive caretaker-primate interaction |
| Does **not** assess any **behaviour** of interest | Does not assess behaviour<br>Assess other behavioural outcomes not fitting the categories of interest. |
| Studies **without** a group/time point that serves as **a control** | Descriptive studies |

### Study details and outcome data extraction

**Supplementary table 2.** Methodological details and outcomes extracted from each unique experimental comparison.

| **Methodological details** | |
|---|---|
| **Study ID** | ✓ Title<br>✓ Corresponding author and e-mail address<br>✓ Year |
| **Subject characteristics** | ✓ Experiment/design<br>✓ Experimental groups<br>✓ Group size<br>✓ Species (*Macaca fascicularis, Macaca mulatta, Macaca arctoides, Macaca nemestrina*)<br>✓ Age<br>✓ Sex (F, M, mixed)<br>✓ Rearing:<br> o Wild-born<br> o In groups |

| | |
|---|---|
| | o Indoor mother-reared<br>o Nursery reared<br>o Mixed |
| **Intervention characteristics for control and exposed groups** | ✓ Country where the primate facility is located<br>✓ Type of facility:<br>o Research facility (academia/public institution)<br>o Research facility (company/private institution)<br>o Sanctuary<br>o Rescue center<br>o Zoo<br>o Other<br>✓ Type of housing:<br>o Single-housing<br>o Protected contact housing<br>o Intermittent contact pair-housing<br>o Pair-housing<br>o Group-housing (> 2)<br>✓ Location (indoors, outdoors, mixed)<br>✓ Cage size<br>✓ Animals per cage<br>✓ Enrichment in control: enrichment in the home cage before the enrichment strategy of interest was introduced or the enrichment strategy for the control group.<br>o None<br>o Physical enrichment (cage size increase, furniture [e.g. perches, visual barriers, porches/verandas/balconies], manipulable objects)<br>o Feeding enrichment (limited to objects incorporated in the home cage that promote foraging [foraging devices])<br>o Mixed (incorporating both types of enrichment)<br>✓ Description of enrichment in control<br>✓ Type of enrichment introduced:<br>o Physical enrichment<br>o Feeding enrichment<br>o Mixed<br>✓ Description of the enrichment introduced<br>✓ Length of enrichment |
| **Outcomes** | ✓ Positive welfare-related behaviors:<br>o Affiliative<br>o Foraging<br>o Infant care and handling<br>o Locomotion<br>o Object manipulation/exploration<br>o Sexual<br>✓ Negative welfare-related behaviors:<br>o Aggression<br>o Harmful self-directed<br>o Inactivity<br>o Stereotypies<br>o Submissive/fearful<br>o Abnormal: including behaviors belonging to the categories of harmful self-directed behaviors and stereotypies that were reported together and therefore could not be assigned to a single category. It also includes other types of abnormal behaviors such as urine drinking or coprophagy<br><br>The description/definition of each of the behaviors reported in a study was checked, only the behaviors that fitted into the categories of interest were extracted along with its definition. |

The authors of 46 studies from which we had incomplete information were contacted by e-mail in hope of having access to the complete dataset or the missing data. If no response was received within a week, and after exhaustion of other alternatives for contacting the authors or completing the data, the study was excluded from the meta-analysis. These studies were still included in our paper in a descriptive analysis.

When a study reported a single behavioral outcome combining behaviors belonging to a mix of behavioural categories, we applied the following rules for extraction: if the outcome contained a mix of positive or negative behaviors from different categories, we extracted the dominant category, and the second most frequent was extracted as an *alternative* category; outcomes combining neutral behaviors (those not of direct interest) with positive or negative behaviors, or combining both positive and negative behaviors, were not extracted.

Examples:

1. Mix of positive behaviors:

   Play: Amicable behavior directed towards another individual that includes play face, play chasing, play biting and non-social play, e.g. object manipulation.

   Behavioral categories: affiliative (dominant category), Object manipulation and exploration (alternative category)

2. Mix of negative behaviors:

   Aggressive/fearful: a discrete aggressive or fearful action or gesture directed towards the cage mate (barking/grunting, hitting/pushing, fear grimacing, biting).

   Behavioral categories: Aggression (dominant category), Submissive/fearful (alternative category)

These alternative categories were used in a sensitivity analysis of the descriptive portion of our study, see “Descriptive analysis”.

### Results

The full list of R packages used in the study is reported separately for each type of analysis at the beginning of their respective R script.

#### List of included studies

37. Jensen, G. D., Bobbitt, R. A., & Gordon, B. N. (1968). Effects of environment on the relationship between mother and infant pigtailed monkeys (Macaca nemestrina). *Journal of comparative and physiological psychology*, *66*(2), 259.

60. Paterson, E. A., O'Malley, C. I., & Turner, P. V. (2024). Sleep quality in cynomolgus macaques (Macaca fascicularis) varies by housing type and following surgery. *Applied Animal Behaviour Science, 272,* 106188.

#### Characteristics of the included studies

**Datasets:** *Full_StudyDetails.xlsx, Densities.xlsx*

**Data script:** *Study_details.html*

The study details of the individual experiments are reported in the dataset: *Full_StudyDetails.xlsx*

The following are brief descriptions of all variables in the dataset:

| | |
|---|---|
| **Study_ID** | A unique identifier for each individual experiment included. |
| **Year** | The year in which the study was published. |
| **Title** | The title of the research paper. |
| **Country** | The country where the research took place. |
| **Facility** | Type of facility where the macaques were housed. |
| **Species** | The species of the macaque used in the experiment |
| **Design** | The design of the experiment (WS, within subjects; CE, control vs exposed) |
| **Sex_C; Age_C; Rearing_C** | Sex/Age/rearing conditions of the animals in the control group/baseline |
| **Sex_E; Age_E; Rearing_E** | Sex/Age/rearing conditions of the animals in the exposed group.<br>For WS experiments, it is NA by default |
| **Housing_C** | The type of housing conditions for the animals in the control group / Baseline phase |
| **CageL_C** | The location of the home cage in the control group / Baseline phase |
| **Enrichment_C** | The type of environmental enrichment provided to the control group / Baseline phase |
| **Description_EC** | The description of the enrichment used in the control group / Baseline phase |
| **CC1_H; CC1_W; CC1_L** | The height (H), width (W), and length (L) in meters of the home cage in the control group / Baseline phase |
| **CC1_floorA** | The floor area of the home cage in $m^2$ in the control group / Baseline phase |
| **CC1_Animals** | The number of animals housed in the home cage in the control group / Baseline phase |
| **CC1_animals/m2** | The density (animals per square meter) in the home cage in the control group / Baseline phase |
| **CC1_m2/animal** | The individual space allowance (square meter per animal) in the home cage in the control group / Baseline phase |
| **CC2_...; CC3_...** | The dimensions, density and space allowance of additional home cages used to house the animals in the control group / Baseline phase (if different from cage 1) |
| **Housing_E** | The type of housing conditions for the animals in the exposed group / Intervention phase |
| **CageL_E** | The location of the home cage in the exposed group / Intervention phase |

| | |
|---|---|
| **Enrichment_E** | The type of environmental enrichment provided to the exposed group / Intervention phase |
| **E_type** | The subtypes of physical enrichment provided to the exposed group / Intervention phase |
| **Description_EE** | The description of the enrichment used in the exposed group / Intervention phase |
| **Length_E** | The length of the enrichment intervention tested |
| **EC1_H; EC1_W; EC1_L** | The height (H), width (W), and length (L) in meters of the home cage in the exposed group / Intervention phase |
| **EC1_floorA** | The floor area of the home cage in $m^2$ in the exposed group / Intervention phase |
| **EC1_Animals** | The number of animals housed in the home cage in the exposed group / Intervention phase |
| **EC1_animals/m2** | The density (animals per square meter) in the home cage in the exposed group / Intervention phase |
| **EC1_m2/animal** | The individual space allowance (square meter per animal) in the home cage in the exposed group / Intervention phase |
| **EC2_...; EC3_...** | The dimensions, density and space allowance of additional home cages used to house the animals in the exposed group / Intervention phase (if different from cage 1) |

#### Characteristics of the two types of experiments

Over half of the experiments (52.6%) that had a **within-subjects design** used experimental groups that were mixed in terms of sex, while a quarter employed groups composed of either males or females (**Figure S5**). Information about the early rearing environment (the first six months) was missing for 59.2% of these experiments. In 18.4% within-subjects experiments, the experimental group had subjects with mixed rearing conditions. In the rest of the experiments, the subjects were either raised in groups (10.5%); born in the wild (6.6%), raised with their (foster) mother only (2.6%), or in a nursery (2.6%) (**Figure S5**).
Of the experiments having a **control vs exposed design** (n = 26), 76.9% used groups that had a mix of both male and female subjects. Females only were used in 11.5% of these experiments, while male-only constellations made up 7.7%. In one experiment, the sex of the subjects was not reported. Control and exposed groups differed slightly in the distribution of their rearing conditions (**Figure S5**). This difference was driven by 6 experiments (single study) in which the control groups had mixed rearing conditions, or unknown (only one experiment), while the exposed groups were raised in groups. In the rest of the experiments, both groups had the same rearing conditions: 7.7% were wild-born or nursery reared (2 experiments each), one experiment employed macaques raised with their (foster) mother only, and in 23.1% of experiments, the rearing conditions of both groups was unknown.

**Figure S6** illustrates the types of enrichment present at baseline/control and during the intervention phases, categorized by experimental design. Within-subjects experiments exhibit a similar pattern to the analysis of all experiments combined (**Figure 2**). In contrast, control-exposed experiments either lacked any enrichment at baseline or employed physical enrichment.

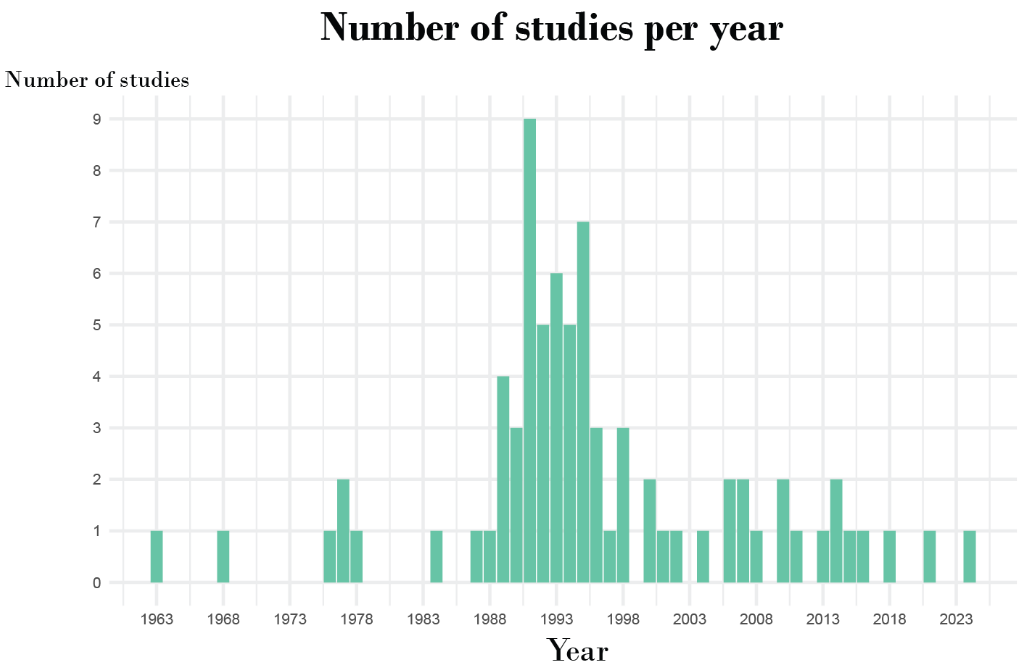


**Figure S3. Studies per year.** The included studies were published between 1963 and 2024 with the majority being published in the 90s.

#### Densities:

The density in control and enriched conditions were calculated using the floor area of the cages (reported or calculated from the cage dimensions) and the number of animals housed in the cages. Whenever the studies reported a range in the number of animals, the highest number was used in the calculation. Whenever the floor area of the cages was reported in ranges, the smallest floor area was used. 17 experiments did not have complete data to perform the calculation of the densities, so calculations of median and mean densities were based on 85 experiments. Animal density calculations did not account for individual animal size; infants or adults were considered equal in contributing to the "animals per square meter" metric.

We opted to calculate animal density based on square meters rather than cubic meters. This approach was chosen due to the higher prevalence of missing height data for cages and to ensure a consistent, meaningful metric across both indoor and outdoor housing environments. For outdoor housing without roof, cubic meter measurements would not be applicable.

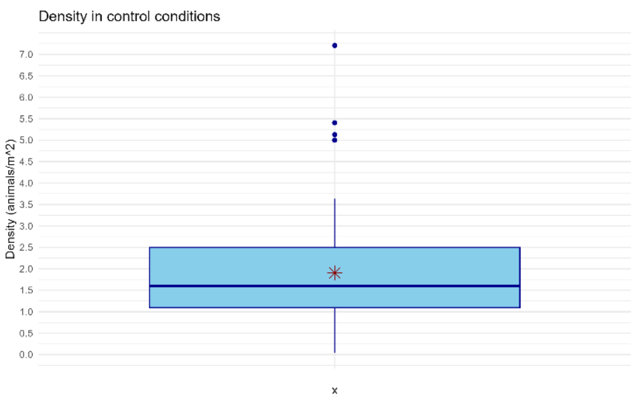


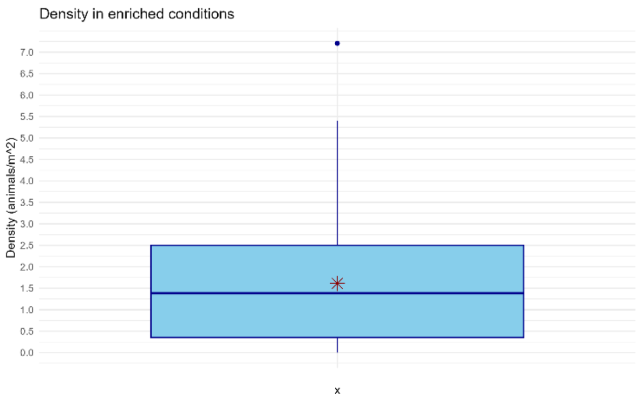

The density of animals in control conditions varied, with the majority (middle 50%) of cages having between 1.1 and 2.5 animals per square meter. The typical density (median) was about 1.6 animals per square meter, and the average density was slightly higher at approximately 1.9 animals per square meter. The densities ranged from a minimum of about 0.4 to a maximum of about 7.2 animals per square meter (adults and infants housed together).
The density in the enriched conditions was slightly smaller than in the control conditions and with wider spread. The majority of cages had between 0.4 and 2.5 animals per square meter. The typical density (median) was about 1.4 animals per square meter (average density was slightly higher at 1.6 animals per square meter). The densities ranged from a minimum of about 0.0002 to a maximum of about 7,2 animals per square meter

For enhanced clarity and direct interpretability, we also calculated the individual space allowance (in square meters per animal).

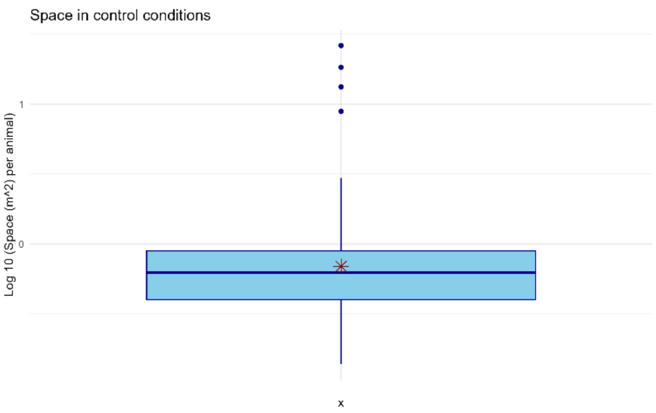


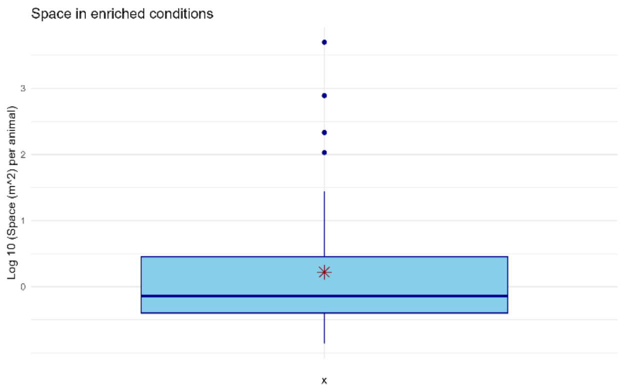


The data for space was log-transformed to normalize its distribution and facilitate clearer visualization, particularly given its wide range and very large outliers.

### Data synthesis

#### Meta-analysis

**Datasets:** *Meta_analysis_Data_R.xlsx, Scatterplot.xlsx* (used for plotting the effects seen in Fig.3), Scatterplot_subtypes.xlsx (used for plotting the effects seen in Fig.4)

**Data script:** *Meta_analysis.html*

This script shows all the results discussed here and in the main article.

For a complete description of the variables in the dataset, please refer to the explanation above, in the section "Characteristics of the included studies". The following are additional variables specific to this dataset:

| | |
|---|---|
| **Study** | A unique identifier for each STUDY included. |
| **Experiment_ID** | A unique identifier for each individual EXPERIMENT included. |
| **Behavior** | The type of behavior (positive or negative welfare-related) |
| **Beh_cat** | The behavioral category of the behavior |
| **Alt_Beh_cat** | The alternative behavioral category, if relevant. |
| **Name** | The name of the behavior as given by the study from which it was extracted |
| **Definition** | The description of the behavior given by the study from which it was extracted |

| | |
|---|---|
| **Extracted_from** | Where in the manuscript the data was extracted from |
| **Unit** | The unit of the outcome |
| **Nc** | The number of animals in the control group / Baseline phase |
| **Mc** | The means of the control group/baseline |
| **Sc** | The standard deviation of the control group/baseline |
| **Ne** | The number of animals in the exposed group / Intervention phase |
| **Me** | The means of the exposed group / Intervention phase |
| **Se** | The standard deviation of the exposed group / Intervention phase |
| **Notes** | Relevant notes |

The multilevel meta-analysis was run with the rma.mv function from the *metafor* R package. For inference about the pooled effect estimates, we used an approach that addresses the analytical complexities of multilevel meta-analysis where conventional Knapp-Hartung adjustments are not directly applicable (test= "t" and dfs="contain" in the *metafor* R package)[5]. We conducted a three-level meta-analysis with robust variance estimation (RVE) for structural enrichment as a whole and for each of the three major types of enrichment (physical, feeding, mixed). A multilevel meta-analysis with RVE handles dependent effect sizes without requiring to explicitly specify the correlation between behaviors, which was unknown[6]. Robust standard errors and confidence intervals for all model coefficients were obtained using the clubSandwich R package, with clustering performed at the study level to account for all within-study dependencies. The forest plots of each analysis are shown in **figure S7**. Pooling and weighting of individual effect sizes (standardized mean differences, SMD - Hedges' g) was done with a random-effects model using the restricted maximum likelihood (REML) for estimating $\tau^2$ at level 2 (variability between different experiments *within* the same study) and at level 3 (variability between studies). The statistical measure for assessing heterogeneity was $I^2$. Two heterogeneity measures were calculated, each quantifying the percentage of total variation associated with either level 2 or 3 ($I^2$ level 2, and $I^2$ level 3, respectively).

The effect of the enrichment interventions was evaluated by comparing pre and post-enrichment behavior and by comparing behavior in control and exposed animals. In the former case, the group size was split in two to avoid double-counting[7].

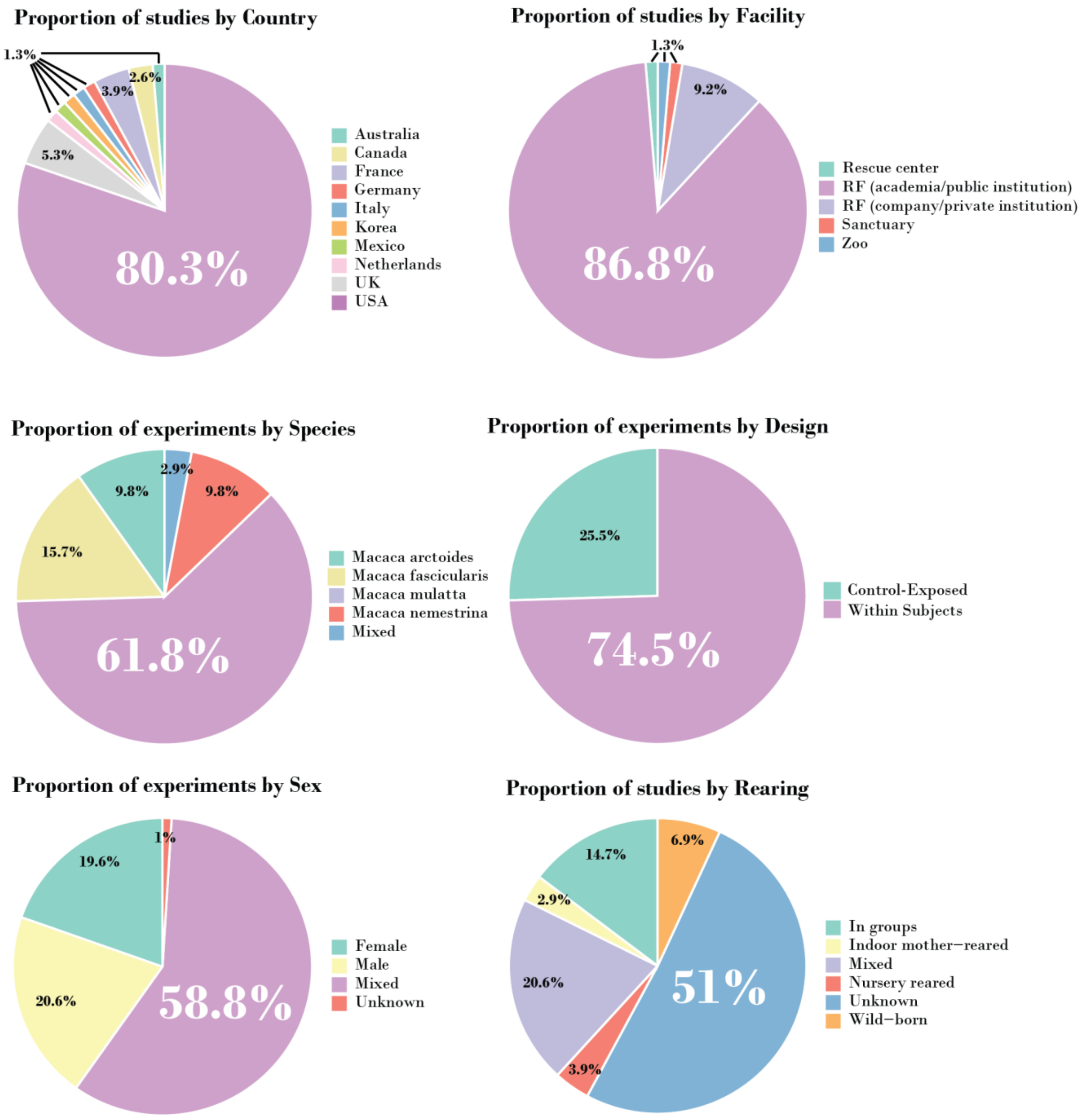


**Figure S4. Characteristics of the included studies.** Most studies were published by research groups based in the USA and conducted at research facilities in academic or public institutions. 61.8% of the experiments were in Rhesus macaques, followed by Long-tailed macaques. The majority of the experiments had a within-subjects design, only 25.5 % used a control versus exposed design. Almost 60% experiments used groups composed of both females and males, while the rest was distributed equally for males and females. In 51% of the experiments, the rearing conditions of the animals was not reported. Abbreviations: RF: Research facility.

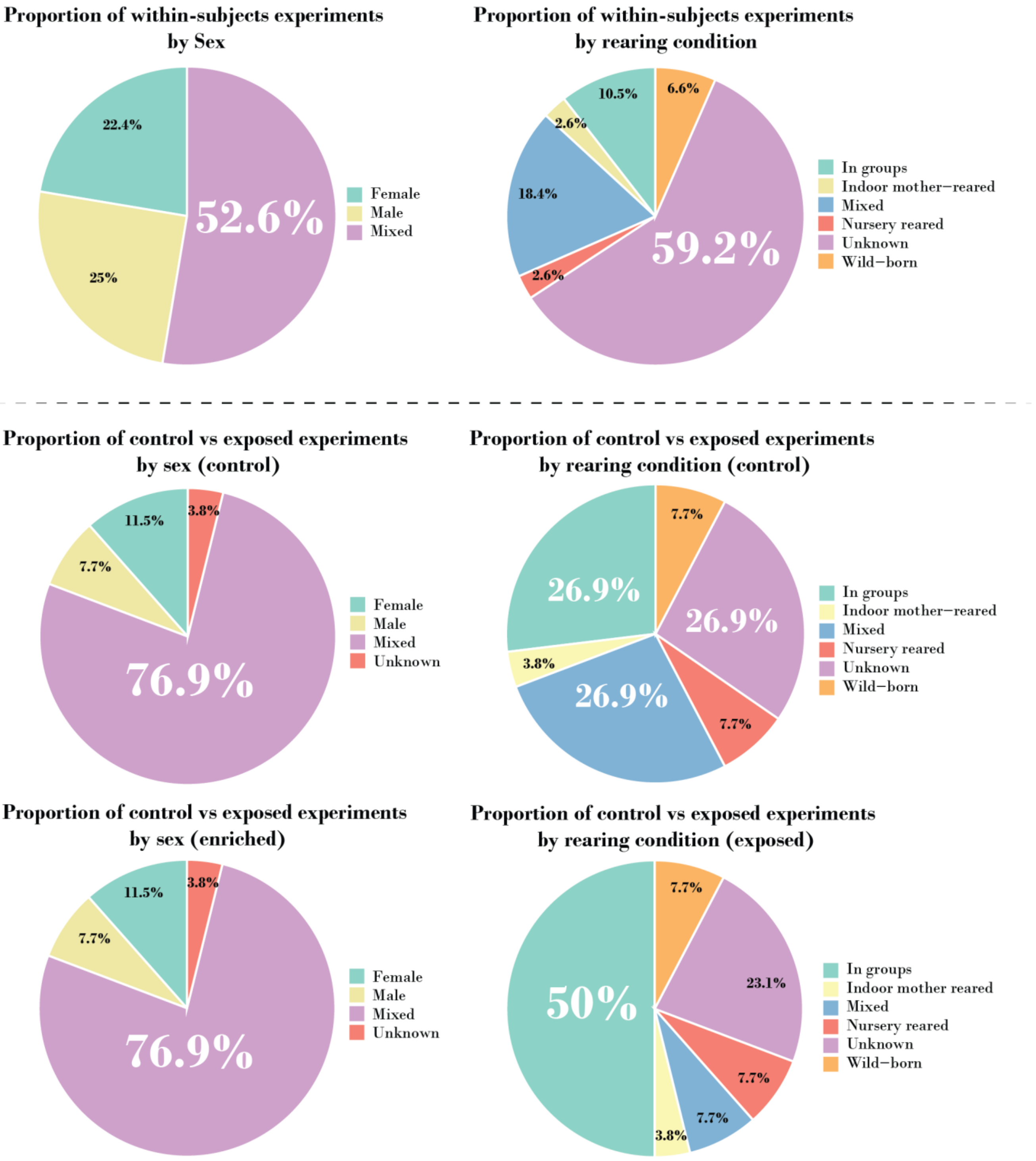


**Figure S5. Characteristics of the included studies split by study design.** Upper part of the figure shows the distribution of within-subjects experiments by sex and rearing condition. The bottom part does the same for control vs exposed experiments but further differentiated by experimental groups. Rearing Condition was defined as the type of environment where the animals spent most of their time during the first six months of their life. This could be: **In groups**: corrals, sheltered housing, environmentally controlled enclosures, and indoor groups (including when reared with the mother in groups); **Indoor mother reared**: indoor caging raised with biological or foster mother only; **Nursery reared**: raised in indoor nursery without monkey's mother, can be paired with conspecifics; **Mixed**: either different animals in the group had different rearing conditions, or the subjects were exposed to different environments during the first 6 months of life; and **Wild-born**: when born in a natural habitat.

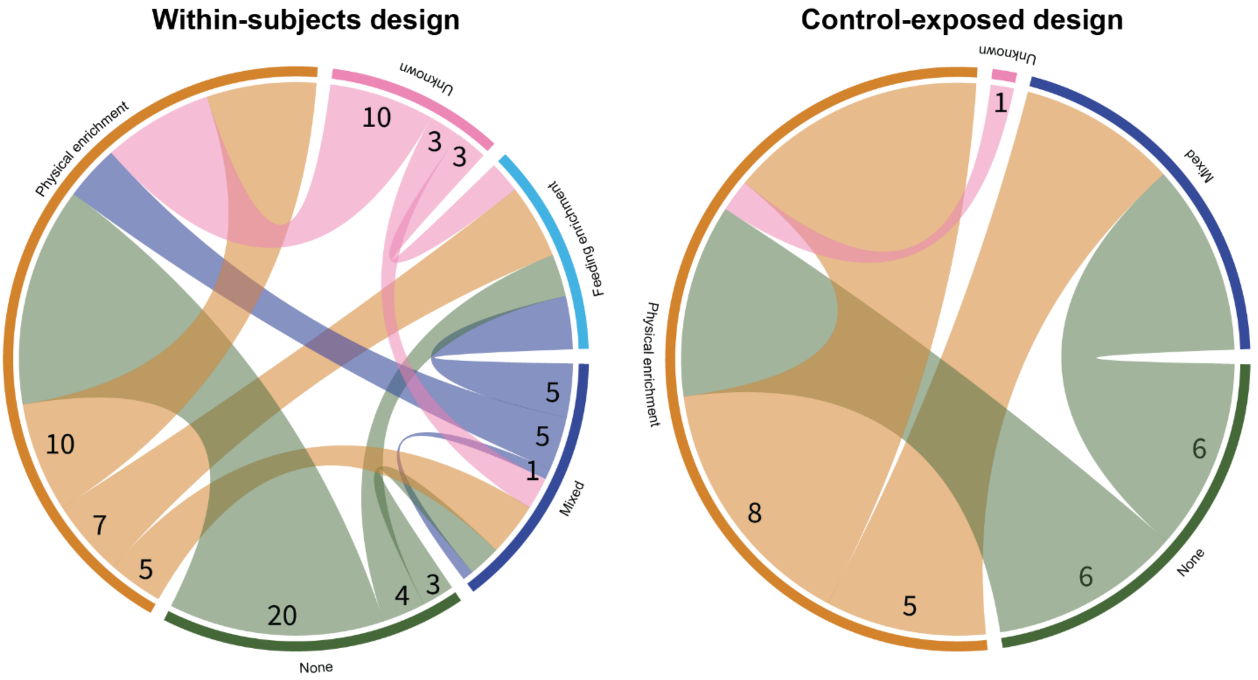


**Figure S6. Enrichment at baseline/control and at intervention phases for the two types of experiments.** This chord diagram shows the number of experiments based on the type of enrichment present at baseline/control conditions and the type of structural enrichment intervention tested. The outer ring segments represent the types of structural enrichment (Physical, Feeding, Mixed, None, or Unknown), with the numbers indicating the start of the connection and representing the total count of experiments starting with that condition. The chords connecting these segments end at the intervention types tested in these experiments. For example, among the within-subjects design experiments with no baseline/control enrichment (None), 20 investigated the effect of introducing physical enrichment, but only 6 of those having a control-exposed design did so.

#### Modified data

The data of four experiments needed to be modified to calculate their effect sizes:

1. Erwin_1976_C2: For this experiment, the unit of analysis was the group, not the individual members. Only two groups were compared with a within-subjects design, meaning that with our adjustment for sample size, the n was reduced to 1 which precludes the calculation of the effect sizes for 9 behaviors. Instead of discarding the data we decided to increase the sample size to 2.
2. Lam_1991_C1 & Lam_1991_C2: The same for these two. The total n is 3 but, because of a within-subjects design, the n was adjusted to 1.5 and the effect size could not be calculated. The group size was increased to 2.
3. Champoux_1990_ Self clasp: the mean and standard deviation (SD) in the enriched group were zero, and the SD of the control group was also zero, making the estimate for the pooled variance zero which prevents the calculation of the effect size. We decided to increase the SD of the control group to 0.049 because this is the biggest effect size we can have that rounds up to 0.0 as reported in the study[8].

They all reported on negative behaviors. We performed a **sensitivity analysis** removing these modified data and the direction and magnitude of the effect on negative-welfare-related behaviors was not significantly changed, structural enrichment still led to a decrease in negative-welfare-related behaviors (SMD: -0.34; 95% CI: [-0.51, -0.17]; $p = 0.0004$).

##### Outliers and influential studies

Behavior outliers and influential studies (individual studies that have a disproportionately large impact on the overall results of the meta-analysis) were identified and their extracted data verified for data entry errors[9].

To identify behavior **outliers**, we calculated studentized residuals using the rstudent function from the *metafor* package[9]. We considered values with an absolute studentized residual greater than 3 as significant deviations, given that less than 1% of residuals in a normal distribution would exceed this threshold by chance (**Figure S8**). Next, we assessed for **influential cases** that could disproportionately affect the meta-analysis results. We did this at the highest level of aggregation (studies), identifying which *independent contributions* to our body of evidence (the primary studies) have the largest impact on our overall meta-analytic findings. We used the influence function from the *metafor* package and identified influential points as those exceeding three times the mean Cook's D value for our dataset[9]. The "3 × mean Cook's D rule" is a heuristic (a rule of thumb) used to identify these influential cases. It suggests that if a study's Cook's D value is three times larger than the average Cook's D across all studies in our meta-analysis, that study is considered significantly influential. None of the identified studies had Cook's D above 0.5 (**Figure S9**).

Two influential studies (contributing 3 experiments and 3 behavioral outcomes) were identified for positive behaviors, both studies contributing results on foraging behavior. Five studies (contributing 11 experiments and 17 behavioral outcomes) were identified for negative behaviors, the majority (n = 10) related to aggression, and the rest to inactivity, stereotypies and abnormal behaviors.

##### Effects of physical enrichment subtypes

As seen in the table below, the subgroup analysis for physical enrichment subtypes indicated that they did not significantly vary in their ability to explain the heterogeneity observed in our overall analysis (the subgroup analysis was not statistically significant, implying insufficient evidence that the subtypes' effect sizes differ significantly from each other). However, to gain a more granular understanding of each subtype's individual impact on welfare-related behaviors, we proceeded to conduct separate multilevel models for each subtype. **Figure 4** shows the mean effect sizes and their respective confidence regions for each subtype on 'Positive behavior' (representing positive welfare indicators) and 'Negative behavior' (representing negative welfare indicators). From this visualization, we observe distinct patterns among the subtypes, even though the overall subgroup test for heterogeneity was not significant. This highlights that while the *differences between* the subtypes might not be statistically robust enough to explain overall heterogeneity, examining their *individual effects* reveals differential impacts. **Supplementary table 3** summarizes the results of the multilevel model used to create the scatterplot.

**Supplementary table 3.** *Results of multilevel meta-analyses (Robust Estimates) for subtypes of physical enrichment. Extensions were not included, since only three behaviors (1 positive and 2 negative) were included in the analyses.*

| Subtype | K (studies, behaviors) | Estimate | CI, lower limit | CI, upper limit | p-value |
|---|---|---|---|---|---|
| **Positive welfare-related behaviors** | | | | | |
| Space | 4, 7 | -0.0065 | -0.3842 | 0.3713 | 0.9554 |
| Objects | 4, 7 | 0.0267 | -1.1291 | 1.1825 | 0.9434 |
| Visual barriers | 3, 10 | 0.2485 | -0.8967 | 1.3936 | 0.4299 |
| Mixed | 3, 10 | 0.5816 | -0.7898 | 1.9530 | 0.2036 |
| **Negative welfare-related behaviors** | | | | | |

| | | | | | |
|---|---|---|---|---|---|
| Space | 5, 15 | 0.1619 | -0.5898 | 0.9136 | 0.5814 |
| Objects | 7, 21 | -0.2106 | -0.4417 | 0.0205 | 0.0600 |
| Visual barriers | 4, 22 | -0.1423 | -0.6734 | 0.3888 | 0.4179 |
| Mixed | 7, 42 | -0.5454 | -0.9395 | -0.1514 | 0.0223 |

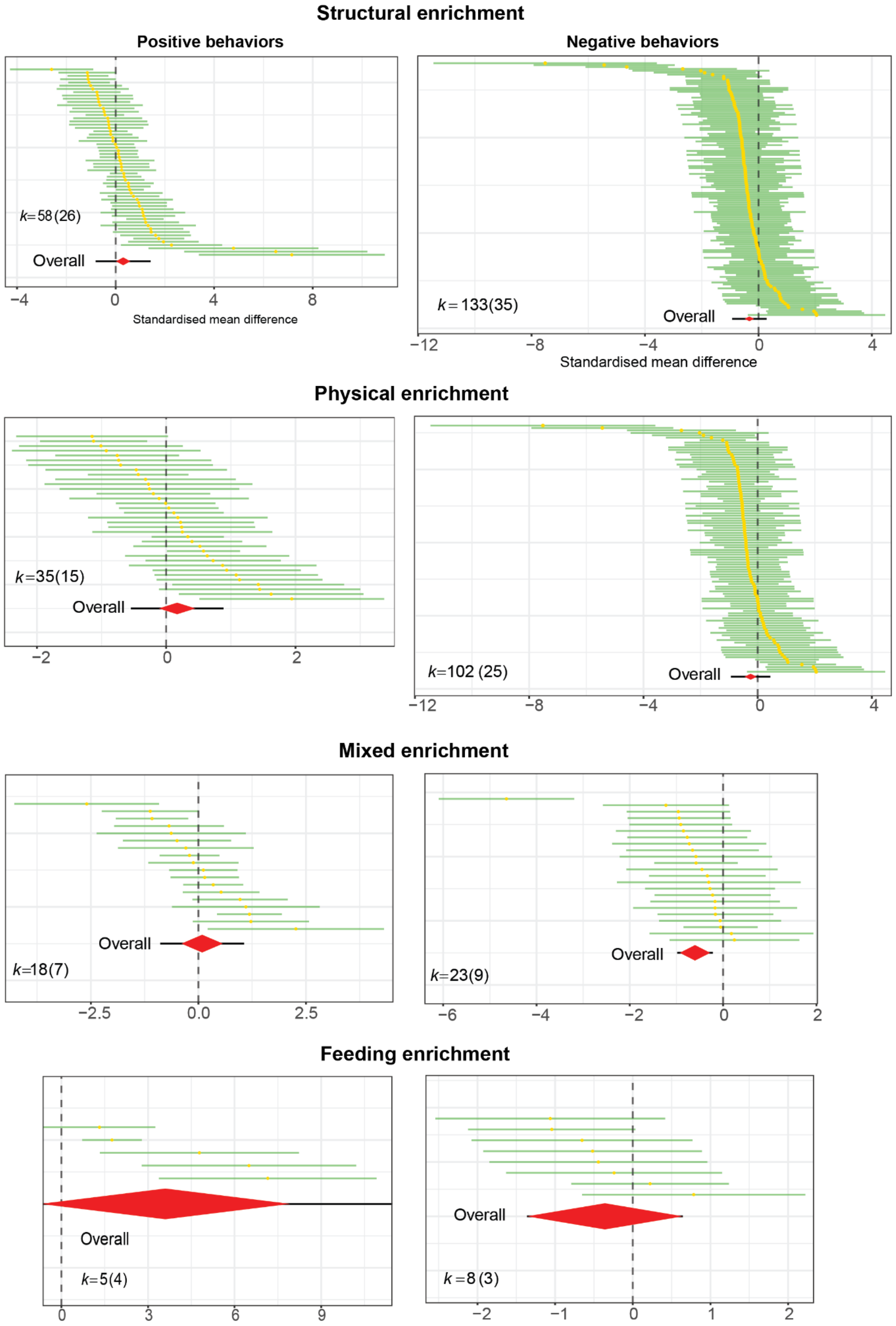


**Figure S7. Forest plots of the types of enrichments (main analyses).** The yellow dots are the effect sizes for each behavior with their confidence intervals (green lines). Red diamonds represent the overall pooled effect sizes. The horizontal width of the rhomboid represents the 95% confidence interval (CI) for this overall pooled effect. The black line is the prediction interval. "k" is the number of behaviors (studies).

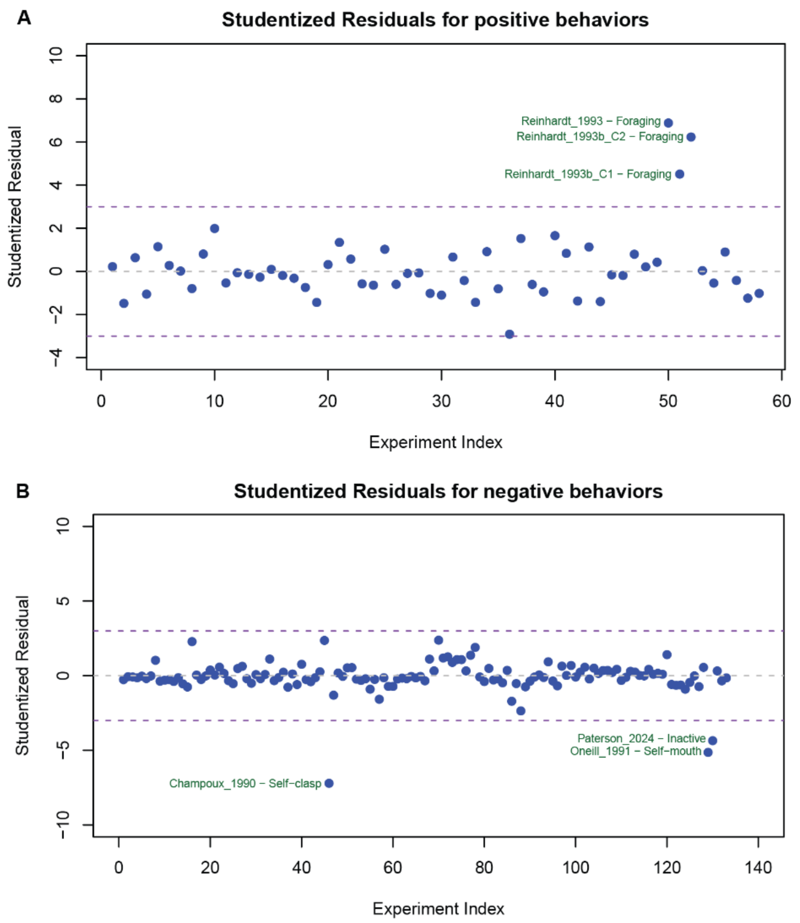


**Figure S8. Outliers.** Behavior outliers were identifying by using studentized residuals for positive (**A**) and negative (**B**) welfare-related behaviors. We checked the outliers for data entry errors and there were none.

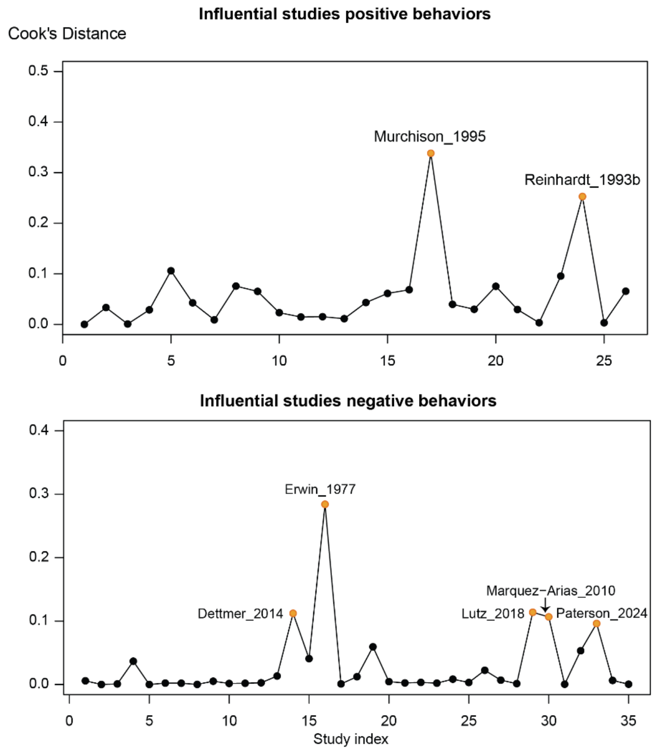


**Figure S9. Influential studies.** The influence of each study in the analysis was calculated using a **"leave-one-out"** approach (re-fitting the meta-analysis model many times, each time omitting one study). Cook's D measures the overall change in the model's fitted estimate when a particular study is removed. Influential studies were those with Cook's D greater than three times the average. Two and five studies were found to be influential in the analysis for positive and negative-welfare-related behaviors, respectively.

**Supplementary table 4.** *Results of subgroup analyses (Robust Estimates).* The subgroups included had a minimum of 2 studies.

| Moderator | K (studies, behaviors)+ | F-statistic (df1, df2) | p-value |
|---|---|---|---|
| **Positive welfare-related behaviors** | | | |
| **Species** (fascicularis, mulatta, nemestrina, and arctoides) | 26, 58 | 3.4126 (3, 1.61) | 0.2754 |
| **Sex_C*** (male, female, mixed) | 25, 54 | 0.5977 (2, 8.16) | 0.5725 |
| **Type of facility#** (Research facility (academia/public institution), Research facility (company/private institution) | 25, 57 | 0.3438 (1, 2.44) | 0.6074 |
| **Housing_C**** (single-housing, pair-housing, group-housing, mixed) | 25, 54 | 0.2278 (3, 1.9) | 0.8714 |
| **Rearing_C*#** (wild-born, nursery-reared, in groups, mixed) | 11, 27 | 9.7805 (3, 0.63) | 0.3504 |
| **Physical enrichment Subtype#** (Space, Visual Barrier, Objects, Mixed) | 14, 34 | 0.8150 (3, 3.87) | 0.5505 |
| *We verified that the values in these variables matched that of the enriched groups for the control vs exposed experiments.<br># **facility:** Only one study, (1experiment, 1 behavior) had a different facility, so it was removed from the analysis.<br>**Rearing**: Only one study (2 experiments, 4 behavior) had indoor mother-reared subjects, so it was removed from the analysis.<br>**Subtype**: Only one study (1experiment, 1 behavior) used extensions, so it was removed from the analysis.<br>** The behaviors in which the housing in the control did not match that of the enriched condition were removed from the analysis (1 study, 2 experiments,4 behaviors).<br>+Unknown values were removed from all analysis whenever present | | | |
| **Negative welfare-related behaviors** | | | |
| **Species#** (fascicularis, mulatta, nemestrina, arctoides) | 34, 132 | 1.1954 (3, 3.05) | 0.4418 |
| **Sex_C*** (male, female, mixed) | 34, 132 | 0.3303 (2, 13.86) | 0.7242 |
| **Type of facility#** (Research facility (academia/public institution), Research facility (company/private institution) | 33, 131 | 1.5832 (1, 2.84) | 0.3018 |
| **Housing_C**** (single-housing, pair-housing, group-housing, mixed) | 34, 116 | 0.2740 (3, 1.19) | 0.8460 |
| **Rearing_C**#** | 14, 46 | 0.2876 (3, 1.56) | 0.8361 |

| | | | |
|---|---|---|---|
| (wild-born, nursery-reared, in groups, mixed) | | | |
| **Rearing_C_ abnormal**# (nursery-reared, in groups, mixed | 10, 26 | 0.2735 (2, 1.6) | 0.7904 |
| **Physical enrichment Subtype**# (Extensions, Space, Visual Barrier, Objects, Mixed) | 25, 102 | 1.2213 (4, 2.7) | 0.4634 |
| *We verified that the values in these variables matched that of the enriched groups for the control vs exposed experiments.<br># **Facility:** Only 2 studies (2, experiments, 2 behaviors) had a different facility (one in a rescue center, and one in a sanctuary), so they were removed from the analysis.<br>**Species:** Only one study (1experiment, 1 behavior) used mixed species, so it was removed from the analysis.<br>**Rearing:** Only 1 study (2 experiments, 17 behaviors) had indoor mother reared, so it was removed from the analysis.<br>**Rearing_abnormal:** Only 1 study (1 experiments, 1 behavior) had wild-born animals, so it was removed from the analysis.<br>** **Housing:** The behaviors in which the housing in the control did not match that of the enriched condition were removed from the analysis (1 study, 2 experiments,17 behaviors).<br>**Rearing:** Removed the behaviors in which the control groups had differing rearing conditions from the enriched group (1 study, 6 experiments, 6 behaviors)<br>+Unknown values were removed from all analysis whenever present | | | |

#### Sensitivity analysis – no enrichment at baseline

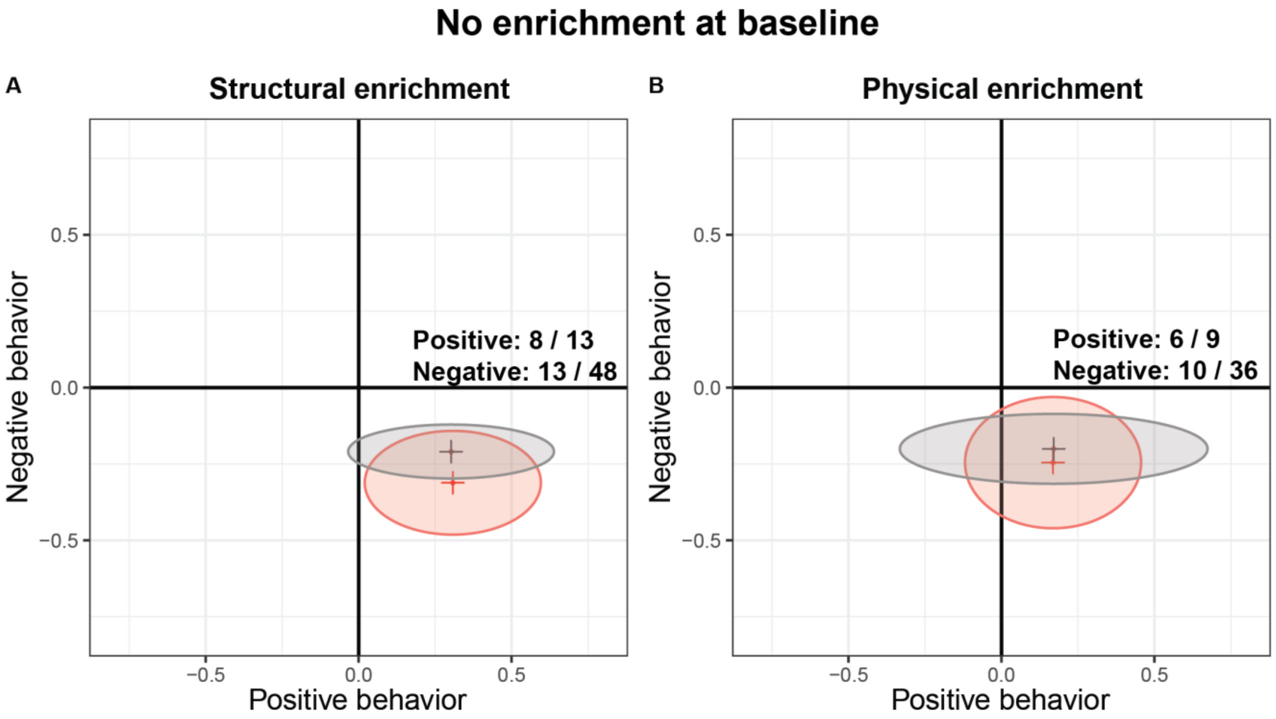


**Figure S10. Sensitivity analysis of studies with no enrichment at baseline.** The X-axis represents the estimated effect on positive welfare-related behaviors, while the Y-axis represents that on negative welfare-related behaviors. The red ellipsis show the effect of the main analysis, while the grey areas show the results of the sensitivity analysis. The number of studies/behaviors pooled for each intervention during the sensitivity analysis are given in the upper right quadrants.

#### Sensitivity analysis – feeding enrichment

We planned to calculate the pooled effect of foraging (structural) enrichment on other behaviors besides foraging, but this was not possible as the studies using this type of enrichment only assessed the effect on foraging behavior.

#### Descriptive analysis

**Datasets:** *Descriptive_R.xlsx*

**Data script:** *Descriptive_analysis.html*

The following are brief descriptions of all variables in the dataset:

| | |
|---|---|
| **Study_ID** | A unique identifier for each individual experiment included. |
| **Design** | The design of the experiment (WS, within subjects; CE, control vs exposed) |
| **Enrichment_C** | The type of environmental enrichment provided to the control group / Baseline phase |
| **Enrichment_E** | The type of environmental enrichment provided to the exposed group / Intervention phase |
| **E_type** | The subtypes of physical enrichment provided to the exposed group / Intervention phase |
| **Behavior** | The type of behavior (positive or negative welfare-related) |
| **Beh_cat** | The behavioral category of the behavior |
| **Alt_Beh_cat** | The alternative behavioral category, if relevant. Otherwise, NA |
| **Name** | The name of the behavior as given by the study from which it was extracted |
| **Definition** | The description of the behavior given by the study from which it was extracted |
| **Reported_result** | The change (increase -decrease) or no difference reported in the study |

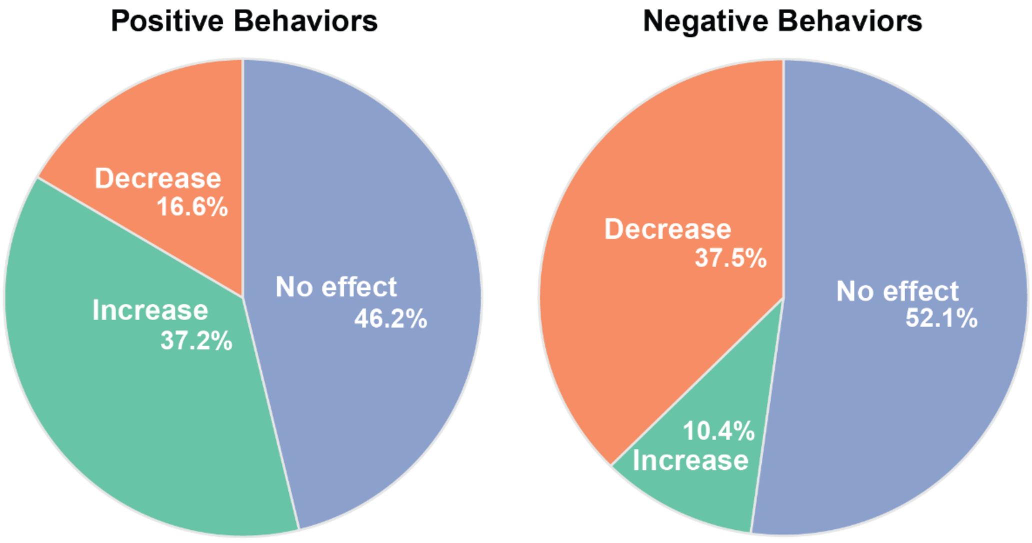


**Figure S11. Distribution of behavioral changes.** Percentage of behaviors that increased, decreased or did not change following a structural enrichment intervention. Positive behaviors (n = 146). Negative behaviors (n = 281).

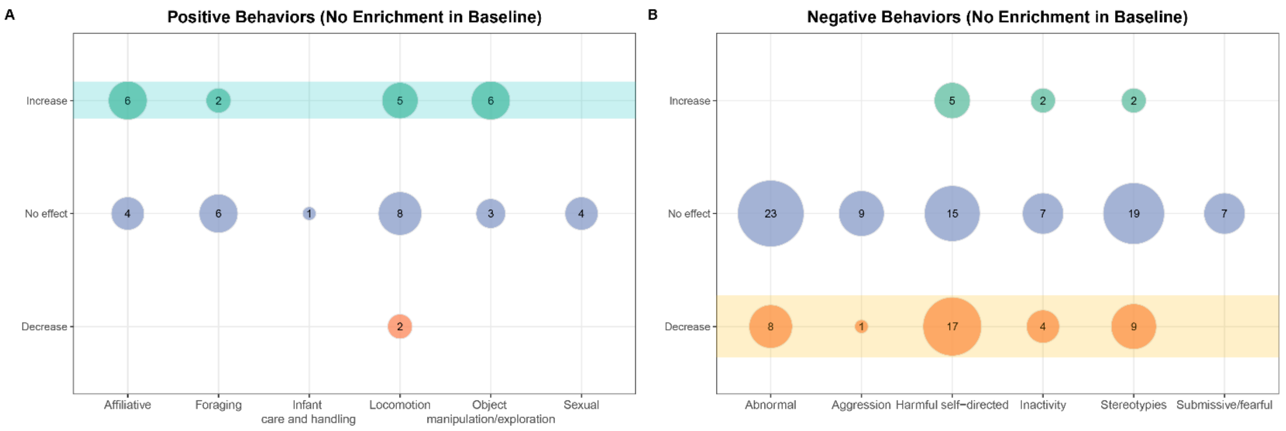


**Figure S12. Bubble plots of behavioral changes for those experiments with no enrichment during the control/baseline condition.** The numbers in the bubbles represent the number of behaviors of each behavioral category that were reported to have increased, decreased or to have not significantly changed after the introduction of structural enrichment in the home cage.

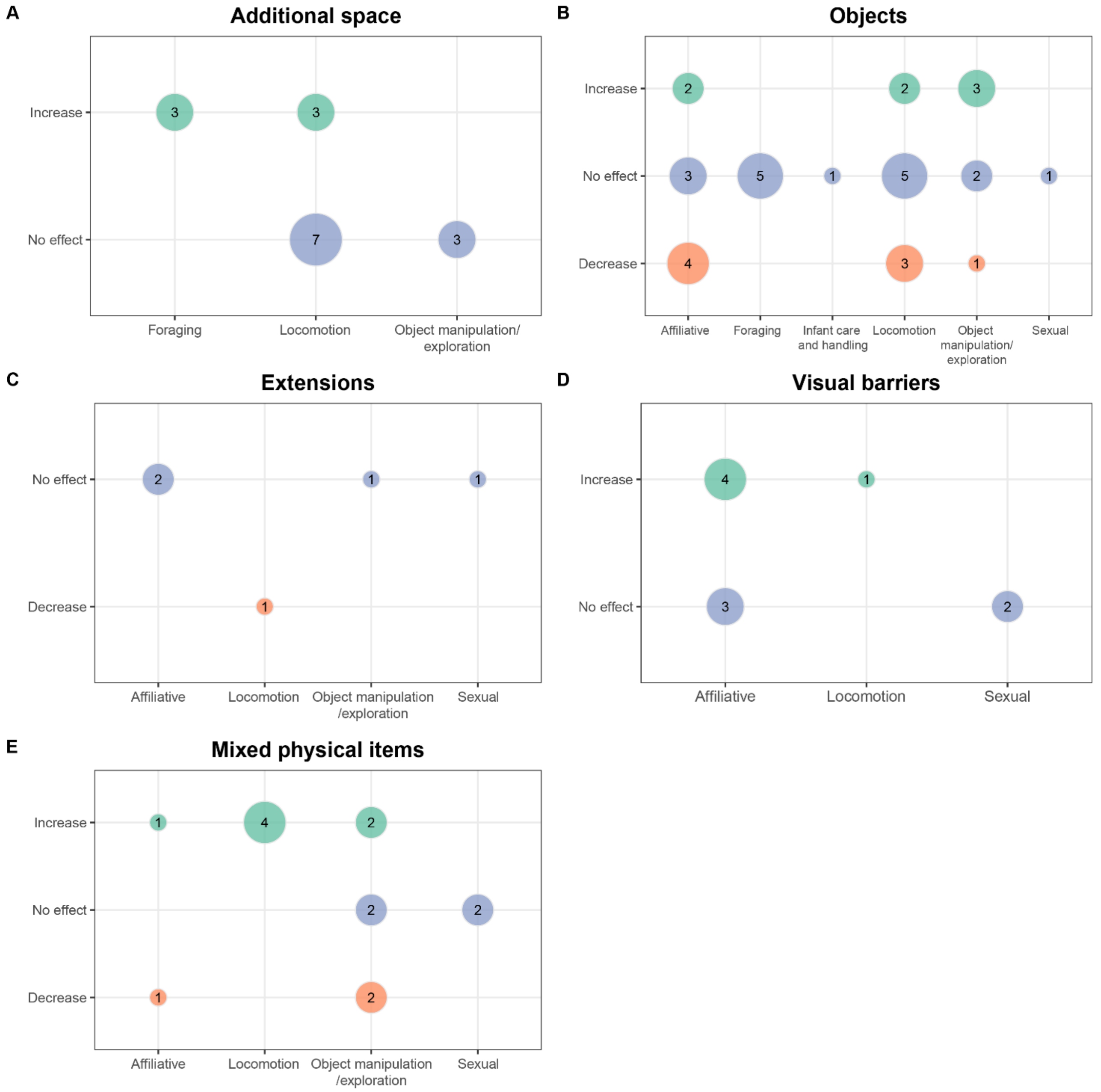


**Figure S13. Bubble plots of changes in positive behaviors for different subtypes of physical enrichment.** The numbers in the bubbles represent the number of behaviors of each positive behavioral category that were reported to have increased, decreased or to have not significantly changed after a subtype of physical enrichment **(A-D)** or a combination of them **(E)** were introduced in the home cage.

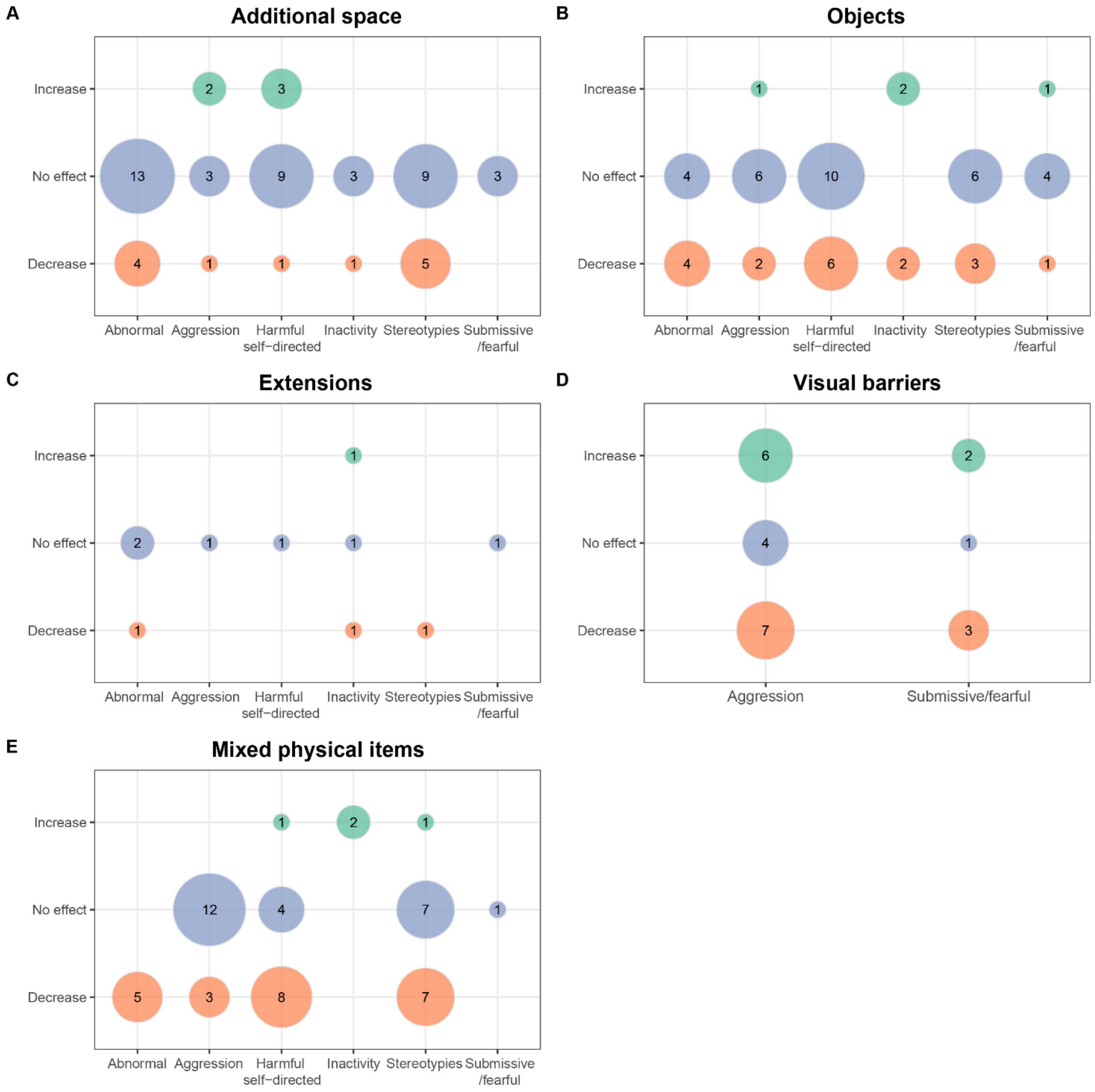


**Figure S14. Bubble plots of changes in negative behaviors for different subtypes of physical enrichment.** The numbers in the bubbles represent the number of behaviors of each negative behavioral category that were reported to have increased, decreased or to have not significantly changed after a subtype of physical enrichment **(A-D)** or a combination of them **(E)** were introduced in the home cage.

A **sensitivity descriptive analysis** was run for structural enrichment using the alternative behavioral categories that were extracted in cases where the behavioral outcome contained a mix of positive or negative behaviors from different categories of interest. There were only 16 instances where the behavioral outcome contained a mix of positive or negative behaviors from different categories. When we used these alternative categories, the overall conclusions drawn from the analysis did not change (**Figure S15**).

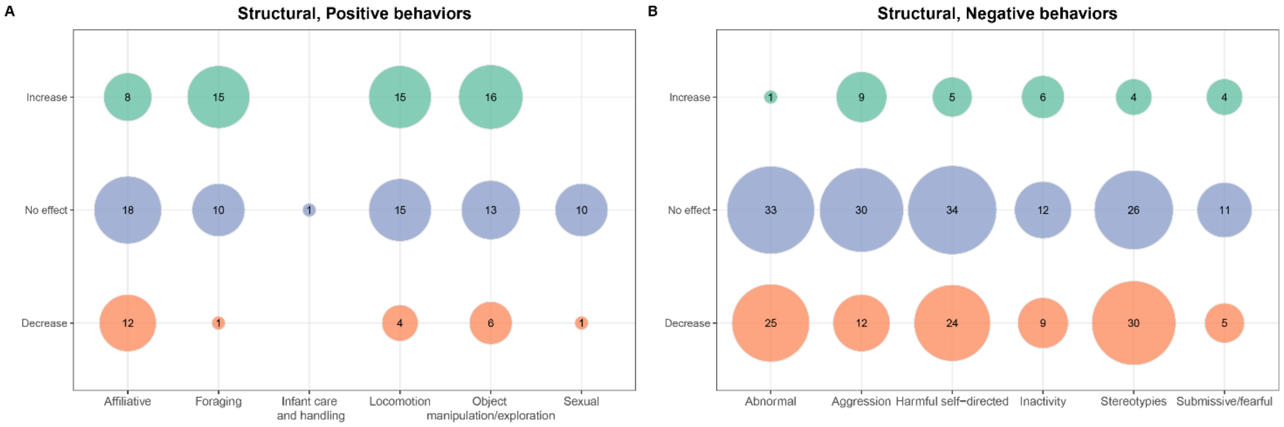


**Figure S15. Bubble plots of behavioral changes using the alternative behavioral categories instead.** There was a total of 16 instances where the behavioral outcome contained a mix of positive or negative behaviors from different categories, the dominant category was extracted as the primary category, while the second most common category as the alternative category. This graph shows the results when the alternative category is used instead. The numbers in the bubbles represent the number of behaviors of each behavioral category that were reported to have increased, decreased or to have not significantly changed after the introduction of structural enrichment in the home cage.
